# Early-life mucosal T cells direct intestinal stem cell fate via a coordinated developmental program

**DOI:** 10.64898/2026.05.08.723752

**Authors:** Madison S. Strine, Kalida Gawon, Long Phan, Weihong Gu, Wenjia Wang, Hanshu Yuan, Dhana Llivichuzhca-Loja, Kerri St Denis, Eduardo G. Gonzalez Santiago, Jiaze Liu, Daniel Zeve, David T. Breault, George Tseng, Liza Konnikova

**Affiliations:** Department of Immunobiology, Yale School of Medicine; Department of Pediatrics, Yale School of Medicine; Division of Animal Sciences, University of Missouri; Bond Life Sciences Center, University of Missouri; Department of Biostatistics, University of Pittsburgh; Renaissance School of Medicine, Stony Brook University; Emory School of Medicine, Emory University; Department of Pharmacology, Yale University; Department of Pediatrics, Boston Children’s Hospital and Harvard Medical School; Department of Obstetrics, Gynecology, and Reproductive Sciences, Yale School of Medicine; Program in Translational Biomedicine, Yale School of Medicine; Program in Human Translational Immunology, Yale School of Medicine; Center for Engineering and Systems Immunology, Yale School of Medicine

## Abstract

Although early-life immunity was once considered immature, the human fetal immune system is dynamic and compartmentalized by the second trimester. By 21 weeks of gestation, T lymphocytes become a major immune population in the fetal small intestine (SI), yet their functional roles within this tissue remain largely undefined. To explore their unique contributions to intestinal development, we established an ex vivo co-culture system in which mucosal T cells isolated from fetal, neonatal, or adult SI donors were cultured with tissue-derived 3D SI organoids derived from various ages. Homeostatic early-life (fetal and neonatal) SI T cells uniquely promoted organoid generation, a metric of stem cell renewal, by upregulating cell cycle-associated gene programs. These early-life T cells also directed intestinal stem cell differentiation toward the secretory lineage in both growth and differentiation phase, highlighting that T cells poise stem cells to adopt secretory fates. T cells from infants with necrotizing enterocolitis (NEC) – an inflammatory intestinal disease affecting predominantly preterm infants – failed to activate these same programs, suggesting a pathologic role for T cells in NEC. T cells from the adult SI similarly failed to support organoid growth or differentiation, revealing developmentally specialized, nonimmune functions for early-life T cells in the intestine. Similarly, T cells derived from cord blood did not enhance organoid generation, indicating that this function is not necessarily a generalized feature of early-life T cells but rather is restricted to mucosal T cells. Organoids derived from adults or NEC, however, could re-enter regenerative states when co-cultured with fetal T cells, indicating that fetal T cells can restore stem cell self-renewal across developmentally and disease-imposed states. We further identified that T cell-derived soluble factors alone were insufficient to modulate intestinal stem cell fate, implying the need for physical interactions. Concordant with this finding, we report that T cells heavily localize to the stem cell niche during prenatal development, where they express factors involved in Notch, Wnt, and growth factor signaling to support fetal stem cell function. Collectively, these findings reveal a coordinated developmental program in which fetal SI T cells balance stem cell self-renewal and differentiation, identifying a developmental immune-epithelial axis that can be harnessed to restore intestinal regeneration.

## INTRODUCTION

The epithelium of the small intestine is maintained by LGR5+ intestinal stem cells (ISCs) in the crypt base^1^. These ISCs give rise to the diverse epithelial cell lineages, including proliferative progenitors (transit amplifying cells), absorptive cells (enterocytes), and secretory cells (Paneth, goblet, enteroendocrine, and tuft cells)^2^. ISCs must balance self-renewal and differentiation to support the functional demands of the intestine, a process that is largely regulated by additional cell types. Within the crypt, ISC identity is reinforced via structural and local signals from other stem cells, Paneth cells, and the subepithelial mesenchyme. Together these cells maintain the crypt as a microenvironment rich in Wingless (Wnt) and bone morphogenetic protein (BMP), among other factors^2,3^. Classically, ISC differentiation in the small intestine requires a gradient-dependent reduction in Wnt signaling as cells transit upward along the crypt-villus axis, with absorptive versus secretory fates being dictated by Notch signaling^2–4^. This process begins in utero, with proliferative ISCs initially being delocalized throughout the intestine. For humans, by 8-12 weeks estimated gestational age (EGA), ISCs already reside in crypts and begin giving rise to terminally differentiated epithelial cells^5^. Enrichment of secretory epithelial cells occurs at approximately 19-20 weeks EGA, with increasing numbers as gestation proceeds^6,7^.

These developmental changes in the epithelial compartment coincide with – and may be functionally linked to – dramatic shifts in the intestinal immune landscape. The human fetal immune system emerges from multiple sites of hematopoiesis only a few weeks into gestation, with adaptive immune cells detectable shortly thereafter^6,8^. Fetal peripheral immune cells traffic to mucosal tissues during the second trimester, including to the intestine^6,8,9^. Though the immune compartment of the small intestine is initially dominated by innate cells, a dynamic shift occurs after 19 weeks EGA^6,10^. Between 19-23 weeks EGA, T cells become the most populous immune cells in the small intestine^6^. Few studies have identified functions for these cells during fetal development, but they co-localize with antigen presenting cells, exhibit T cell receptor signaling, and adopt memory states, suggesting that immune education may begin before birth^10^. Furthermore, fetal intestinal T cells exhibit tissue-residency programs, poising them to directly interact with the epithelium during fetal development^6,10–12^. CD4+ T cells in the fetal intestine are enriched for Th1-like cytokines, like tumor necrosis factor alpha, which has been shown to support ISC proliferation^12^. Notably, T cell programming is dysregulated in preterm infants with necrotizing enterocolitis (NEC), a severe and often fatal gastrointestinal disease characterized by both aberrant immune responses and epithelial barrier dysfunction^6,12,13^. Birth marks a sharp transition in intestinal T cell composition and programming, characterized by a rapid shift toward CD8^⁺^ T cell bias and increased inflammatory and exhaustion signatures^6,14^.

These sharp developmental transitions raise the possibility that intestinal T cells may serve distinct, context-dependent roles before and after birth. Although T cells undergo profound developmental transitions from fetal to adult life, their functional roles across these stages have not been well described. Adult intestinal T cells control infections and modulate tolerance and epithelial function, but the functions these cells play prenatally are less defined^15,16^. Multi-omics studies have offered comprehensive, multi-donor characterization of immune and epithelial cell states during these developmental periods, but functional outcomes of these changes still remain largely undefined^6,7,10,11,14^.

T cells are well positioned to influence epithelial development during gestation, yet the extent to which they directly regulate ISC function across developmental stages remains unknown. Addressing this question has been limited by the lack of experimental systems that allow controlled, parallel comparisons of immune-epithelial interactions across multiple developmental periods. Intestinal organoid models, which can recapitulate the intestinal epithelium, provide an opportunity for reductionist investigation of the developmental cues that shape ISC behavior ^1^. While epithelial organoid models were introduced nearly twenty years ago, the incorporation of other cells comprising the intestinal niche is a more recent development. For example, immune cells have been integrated to organoid cultures, enabling more mechanistic studies of immune-epithelial crosstalk and immune cell function within the intestine^12,17–20^. Here, we introduce a T cell-organoid co-culture model that enables donor-mismatching to systematically interrogate T cell function in the intestine across pre- and post-natal human development. Using this culture system, we identified that early-life (fetal and neonatal) T cells uniquely support intestinal stem cell renewal and secretory lineage differentiation, whereas adult T cells do not. This system also carries disease relevance, as we demonstrated that NEC-associated T cells do not have the same regenerative properties, underscoring a role for T cells in NEC pathogenesis. Excitingly, fetal T cells can induce growth and differentiation of NEC-associated organoids, pointing to a potential therapeutic strategy. We further show that early-life T cells preferentially localize near crypts, where they express factors that support stem cell renewal and differentiation. Taken together, our work reveals that early-life T cells orchestrate a coordinated developmental program shaping intestinal epithelial maturation, with implications for understanding and potentially treating neonatal intestinal diseases.

## RESULTS

### Early-life mucosal T cells promote intestinal stem cell renewal

We generated organoid lines, which model the intestinal epithelium and can be maintained in culture long-term, from cryopreserved human small intestine (SI) tissue (**Figure 1A**). To investigate T cell-epithelial crosstalk, we isolated T cells from these same donors by CD3 positive selection then embedded them in Matrigel^®^ with dissociated organoids (**Figure 1A**). We co-cultured T cells and organoids for six days, allowing organoid formation from intestinal stem cells (ISCs) (**Figure 1A**). To first examine how T cell function may shift during their period of generation and expansion in the fetal SI, we co-cultured organoids and T cells isolated from fetal donors of 19, 21-22, or 23 gestational weeks with no congenital anomalies (**Figures 1B and 1C**). These co-cultures were prepared using both donor-matched and -mismatched combinations of organoids and T cells. Regardless of fetal age or whether T cells were derived from heterologous or autologous tissue, SI T cells significantly increased the number of organoids generated, a metric of stem cell renewal, organoid growth, and differentiation potential (**Figures 1B and 1C**). To see if these properties were conserved postnatally, we then performed co-cultures using organoids and T cells derived from the SI of neonates within the first week of life (**Figures 1D and 1E**). Similar to fetal SI T cells, neonatal SI T cells significantly promoted organoid generation in organoids derived from both fetal and neonatal donors (**Figures 1D and 1E**). This suggests that early neonatal T cells retain regenerative properties, and that both fetal and early postnatal stem cells are responsive to this reprogramming. Not all neonatal T cell donors were able to support organoid generation, and the variation in organoid generation was higher than from fetal SI T cell co-cultures (**Figures 1D and 1E**). We also observed suppression of some cellular proliferation pathways (**Figures S1A and S1B**). Next, we examined whether the tissue source of T cells matters for this phenotype by co-culturing cord blood T cells from term infants with fetal organoids (**Figures 1F and 1G**). In contrast to SI-derived fetal and neonatal T cells, T cell from cord blood were unable to drive organoid generation (**Figures 1F and 1G**). As our data suggested that ISC renewal is driven by tissue resident T cells, we co-cultured fetal or adult organoids with T cells isolated from the SI of adults (20-80 years) to ask whether this crosstalk persists into adulthood (**Figures 1H and 1I**). In stark contrast, to fetal SI T cells, adult SI T cells did not affect organoid generation in fetal or adult organoids (**Figures 1H and 1I**). However, adult ISCs are phenotypically distinct from fetal ISCs^5,21^, leading us to ask whether the adult epithelium could be reprogrammed to this regenerative state by fetal SI T cells (**Figures 1J and 1K**). To answer this question, we co-cultured fetal SI T cells with adult SI organoids and observed substantial organoid generation (**Figures 1J and 1K**). Pathway analysis from these co-cultures indicated activation of cell cycle pathways, underscoring that fetal SI T cells can intrinsically drive ISC renewal by reprogramming ISCs even from aged tissue (**Figures S1C and S1D**). Collectively, these data show that, unlike adult SI T cells, SI T cells from the early-life (fetal and neonatal) human SI uniquely stimulate ISC renewal, enabling organoid formation in an ex vivo co-culture model of T cell-epithelial cell interactions.

**Figure 1.**
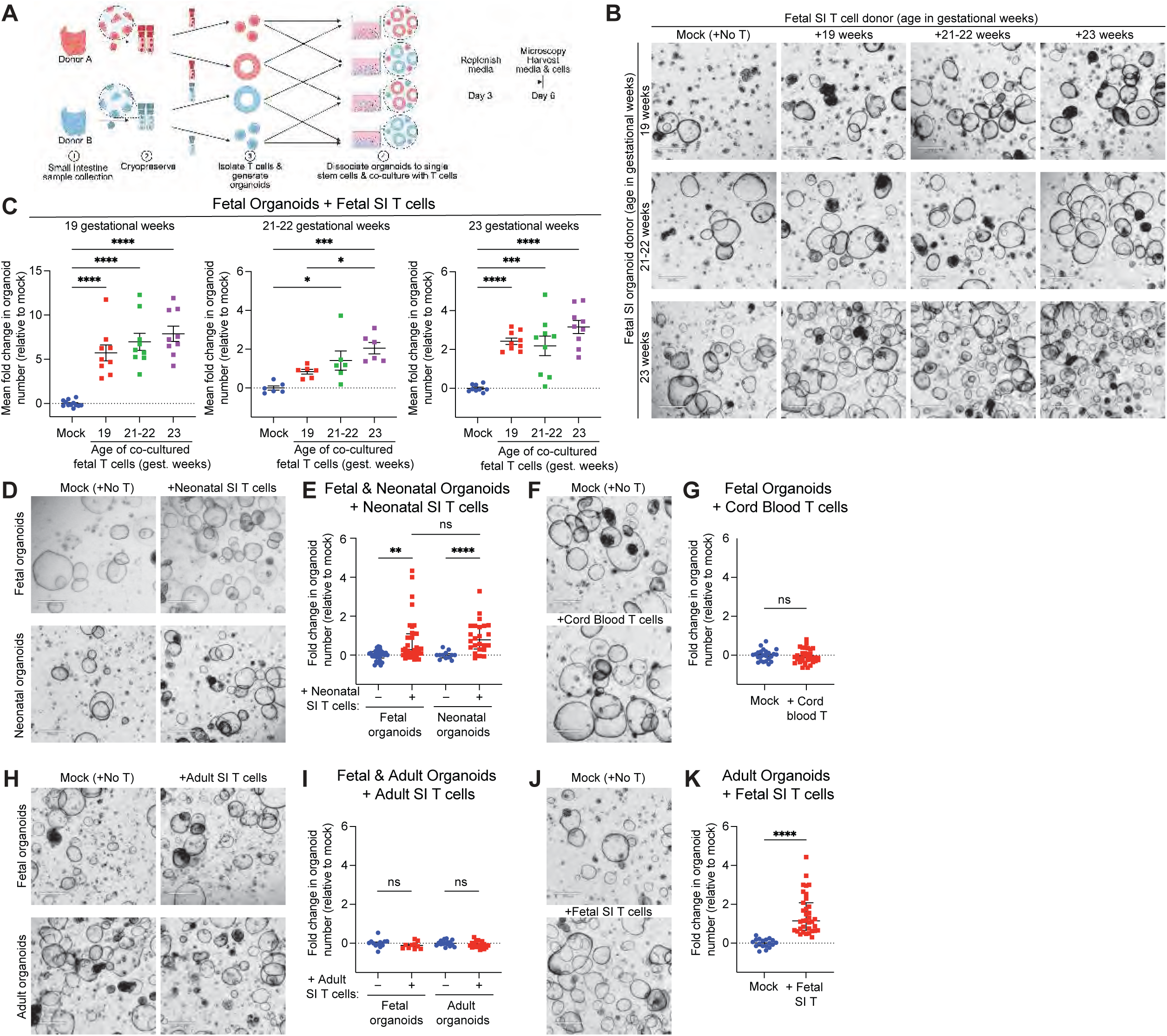
Fetal and neonatal mucosal T cells promote intestinal stem cell renewal. **(A)** Co-culture strategy (created with Biorender.com). Small intestine (SI) was cryopreserved, and stable organoid lines were generated. T cells were isolated from SI tissue **(B-E, H-K)** or cord blood **(F-G)** by positive selection. CD3+ cells were co-cultured with established SI organoid lines generated from fetal **(B-I)**, neonatal **(D-E)**, or adult **(I-K)** donors. **(B, D, F, H)** Representative images from day 6 of co-culture. Scale bar is 530 µm. **(C)** Data were generated from one organoid and T cell donor per fetal age group and analyzed by one-way ANOVA with Tukey’s post-hoc test. Only significant comparisons are denoted. Data are shown as means ± SEM. **(E, G, I)** Data were generated from 2-4 organoid and T cell donors each. Data were analyzed via Mann-Whitney and are shown as medians ± IQR. (K) Data were generated from 3 organoid donors and 4 T cell donors and are normalized to the number of organoids generated in mock (no T cell) conditions, represented as fold change. Each dot represents the average of three technical replicate wells **(C)** or one well of organoids **(E, G, I, K)**. Data were analyzed via Welch’s t-test and are shown as means ± SEM. Each experiment was repeated 2-3 independent times. Each dot represents one well of organoids. Significance is indicated as follows: * p<0.05; ** p<0.01; *** p<0.001; **** p<0.0001; ns, not significant (p>0.05).

### Early-life SI T cells induce expression of epithelial differentiation genes

To investigate whether SI T cells impose additional changes on the epithelium over time, we performed bulk RNA sequencing (bulk RNA-seq) from early (day 3) and late (day 6) time points of fetal SI organoids co-cultured with or without fetal SI T cells (**Figure 2; Figures S2A and SB**). As early as day 3 of culture, organoids demonstrated enrichment in cellular proliferation pathways and negative regulation of epithelial cell apoptosis when grown with fetal SI T cells (**Figures 2A-C**), which would support the organoid formation and growth described in **Figure 1**. Unexpectedly, there was mild enrichment in pathways associated with epithelial differentiation despite these organoids being maintained in growth phase media, including genes associated with secretory cell identity such as *ATOH1* (encodes secretory lineage defining transcription factor), *NEUROG3 (*encodes the enteroendocrine cell (EEC) defining transcription factor), *CHGA, PYY, and SST* (encoding canonical EEC markers), *DEFA6* (encodes a classical Paneth cell-derived antimicrobial peptide), and *SPINK4* (encodes a goblet cell effector involved in mucus structure and repair), among others (**Figures 2A-C**)^16^. By day 6, much of this proliferative programming remained enriched and epithelial cell differentiation showed additional classical goblet cell markers like *MUC5B* and *CLCA1* (**Figures 2D-F**). These data highlight that T cells can skew ISC differentiation along the secretory lineage even in the absence of organoid differentiation media, suggesting this is mediated by factors supplied by T cells. To assess whether fetal SI T cells influence the cellular proportions of differentiated epithelial cells – rather than biasing only their transcriptional profiles – we performed computational deconvolution of these data using a single-cell RNA-sequencing (scRNA-seq) atlas of the pre- and post-natal human SI^6,22^. In concordance with the stem cell renewal phenotype (**Figures 1B and 1C**), we identified an increase in ISCs when fetal SI organoids were co-cultured with fetal T cells (**Figure 2G**). Moreover, co-culture with fetal T cells significantly increased transit amplifying (TA) cells by day 3, which were interestingly mostly lost by day 6, indicating these cells likely differentiated to other fates by this time point (**Figure 2H**). When focusing on the differentiated fraction of epithelial cells, absorptive enterocytes, and EECs did not change in frequency when organoids were co-cultured with fetal T cells (**Figures 2I and 2J**). However, in the presence of fetal T cells, goblet and Paneth-like cells became significantly more frequent (**Figures 2K and 2L**). A similar trend, though not significant, was observed for tuft cells (**Figure 2M**). Together, these data highlight that fetal SI T cells incite gene programming that supports spontaneous ISC differentiation, particularly along the secretory lineage.

**Figure 2.**
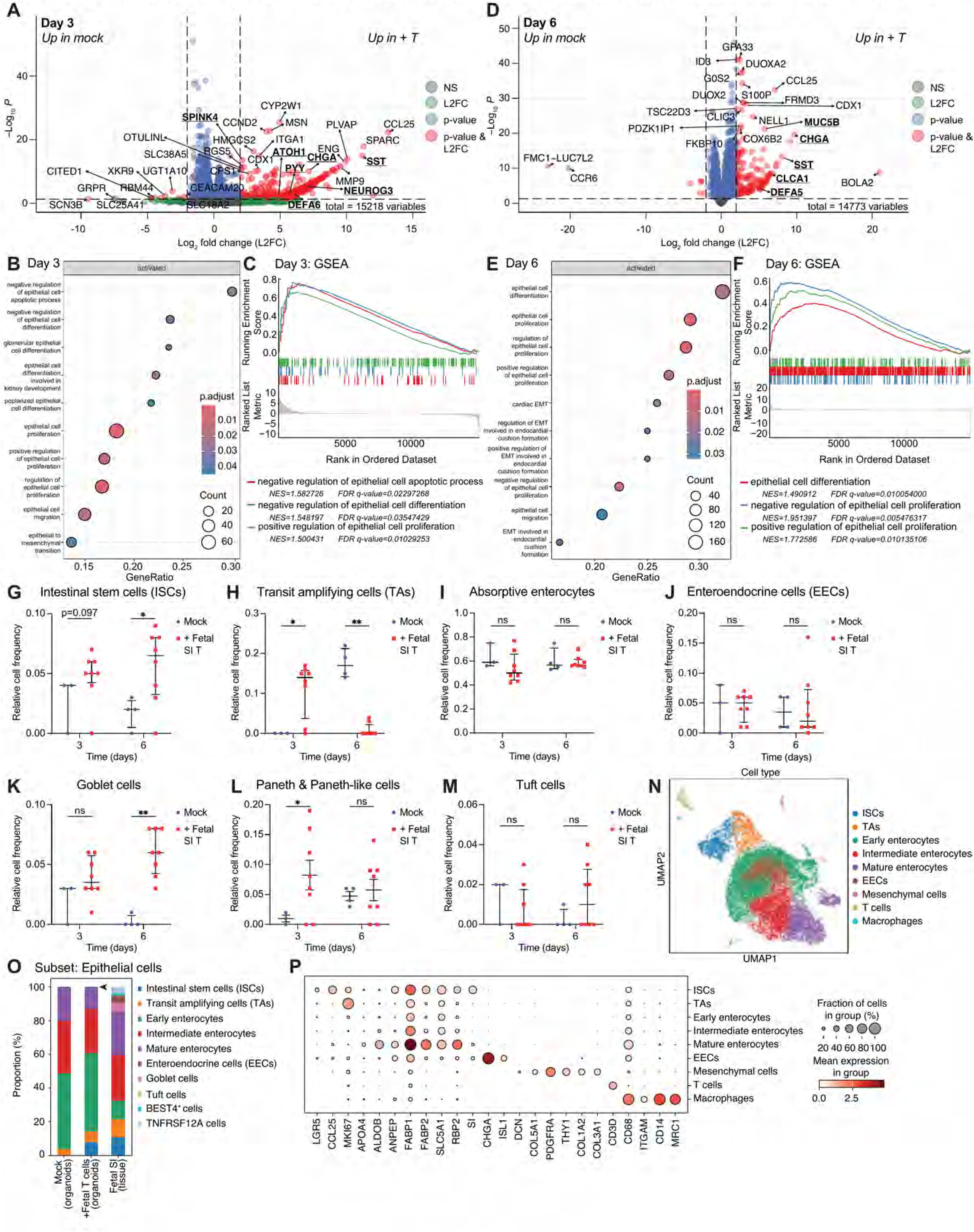
Fetal mucosal T cells drive epithelial proliferation and differentiation in the fetal small intestine. Fetal SI organoids were co-cultured with fetal SI T cells and assessed by RNA-seq at 3 or 6 days of culture in growth phase media. **(A-M)** Bulk RNA-sequencing (RNA-seq). **(A, D)** Volcano plots showing differentially expressed genes (|L2FC|>2, p<0.05) for co-cultures on day 3 **(A)** and day 6 **(D)**. **(B, E)** Gene set enrichment analysis activated intestinal pathways from co-cultures on day 3 **(B)** and day 6 **(E)**. Biological pathways were analyzed via Gene Set Enrichment Analysis (GSEA) Gene Ontology. **(C, F)** Leading edge plots showing enriched pathways for proliferation, survival, and differentiation pathways for co-cultures on day 3 **(C)** and day 6 **(F)**. **(G-M)** Bulk RNA-seq data were computationally deconvoluted to estimate relative cellular frequencies at a pseudo-single cell level for co-cultures at days 3 and 6, showing enrichment for progenitor cells and secretory cells. Each dot represents one sample (pooled 3 technical replicate wells) that were harvested and sequenced. For **(A-M)**, experiments were performed twice with two organoid and T cell donors per sequencing experiment. Data in **(G-K, M)** were analyzed via Mann-Whitney and are represented as medians ± IQR. Data in **(L)** were analyzed via Welch’s t-test and are shown as means ± SEM. Co-cultures collected on day 3 were sequenced via low-input library preparation, precluding direct comparisons to day 6 standard library preparations. Significance is indicated as follows: * p<0.05; ** p<0.01; ns, not significant (p>0.05). (N-P) Single-cell RNA-seq at 3 days of culture. **(N)** Uniform manifold approximation and projection (UMAP) for cell clusters identified. **(O)** Data were subclustered on epithelial cells to identify relative cellular proportions comprising the epithelium of organoids in the presence or absence of T cells, relative to physiologic cell frequencies in the fetal SI ^6^. **(P)** Expression dot plot for a subset differentially expressed genes used in cellular identification and cluster naming.

Although neonatal SI T cells induced some secretory programming (e.g., goblet cell markers *MUC5B* and *MUC4*, the tuft cell transcriptional coactivator *POU2AF3*), signatures for Paneth cells and EECs were missing from co-cultures with neonatal organoids (**Figures S1A and S1B**). Adult SI T cells induced even less secretory identity programming in adult organoids (**Figures S1E and S1F**). *MUC3A* and *MUC13* were upregulated in adult organoids co-cultured with adult SI T cells, but these are transcriptionally more associated with enterocyte identity than goblet cells^16^ (**Figure S1E**). Classical goblet cell genes *MUC5B* and *TFF3*, encoding mucin 5b and a mucus-stabilizing protein, did appear when fetal organoids were co-cultured with adult SI T cells, (**Figure S1G**), but these were absent in adult organoids co-cultured with fetal SI T cells (**Figure S1C**). Aside from these markers, we did not observe a global enrichment in differentiation pathways (**Figure S1H**). These data reveal that the early-life epithelium may have greater propensity to adopt secretory cell fates in general, and fetal SI T cells can better direct ISCs toward secretory differentiated states. To make pairwise comparisons between fetal organoids co-culture with fetal, neonatal, or adult SI T cells, we integrated these datasets (**Figures S2C and S2D**). Fetal SI T cells were better at inducing secretory genes (*CHGA*, *MUC5B*, *DEFA5*, *SST*, etc.) across the board (**Figures S2E and S2F**). As this trend is still detectable but weaker in neonatal organoids and lost in the aged adult epithelium, this may highlight a developmental switch between pre- and post-natal intestinal needs. Overall, these data underscore that both early-life intestinal T and epithelial cells are needed to initiate full secretory lineage programming in the developing gut.

### Fetal SI T cells promote ISC commitment to the secretory lineage

To directly elucidate if fetal SI T cells drive a differentiation program, we performed similar fetal organoid-SI T cell co-cultures, either in growth phase or differentiation phase. Because deconvolution provides only approximate cell type proportions, which may differ from the true cellular composition, we employed scRNA-seq to directly capture individual epithelial cell populations. As early as 3 days of growth phase co-cultures, fetal SI T cells augmented the size of the ISC and TA compartments and led to the emergence of a small number of EECs (**Figures 2N-P**, arrowhead for organoid EECs). In cultures lacking T cells, these secretory populations were completely absent (**Figures 2N-P**). We then applied this scRNA-seq approach to co-cultures maintained in differentiation phase, as terminal ISC differentiation may require additional factors^23^ (**Figure 3**). We first allowed organoid generation for 2 days in growth phase, then substituted the growth media with differentiation media for 4 days, after which we collected cells for sequencing, microscopy, and immunofluorescence (IF) (**Figure 3**). Budding is a proxy for organoid differentiation, caused by an initial increase in TA cell progenitors that can later differentiate into other cell types^1^. With significantly more TA cells at day 3 in the deconvoluted data (**Figure 2**), we quantified bud formation after 4 days of differentiation (**Figure 3**). Organoids co-cultured with fetal SI T cells harbored significantly more buds per organoid with larger buds overall (**Figures 3A-C**). In differentiation phase by day 6, co-culture with fetal SI T cells led to a significant increase in all secretory lineage cells (Paneth cells, goblet cells, EECs, and tuft cells) (**Figures 3D-F**). Interestingly, we also observed a notable increase in intermediate enterocytes (**Figure 3E**). To ensure that the differentiated cells in our organoid cultures were comparable to those found in the tissue, we integrated these organoid data with scRNA-seq from the fetal SI (**Figure 3E; Figure S3A**)^6^. When subsetting on epithelial cells only, fetal organoids more accurately recapitulate the relative cellular proportions of the physiologic fetal SI when co-cultured with T cells (**Figure 3E; Figures S3B-D**)^6^. As transcript detection doesn’t directly equate to protein, to validate these scRNA-seq data, we assessed the presence of chromogranin A (CHGA) and mucin 2 (MUC2), classical protein markers of EECs and goblet cells, respectively (**Figures 3G and 3H**). Organoids co-cultured with T cells showed significantly higher frequency of CHGA- and MUC2-positive cells (**Figures 3G and 3H**). To verify the functionality of these differentiated cells, we measured the abundance of EEC-derived molecules in the supernatant of organoid cultures (**Figure 3I**). In co-cultures, glucose-dependent insulinotropic polypeptide (GIP) was significantly elevated relative to organoids cultured without T cells (**Figure 3I**). Glucagon-like peptide 1 (GLP-1), insulin, and peptide YY (PYY) similarly trended higher in the presence of T cells, though secretin did not (**Figure 3I**). These results suggest that fetal T cells induced functionally competent EECs – most likely K and L cells – given that these subsets predominantly produce GIP, GLP-1, insulin, and PYY, whereas secretin derives from S cells ^24,25^. These findings are overall consistent with more promiscuous peptide secretion from fetal EECs ^24^.

**Figure 3.**
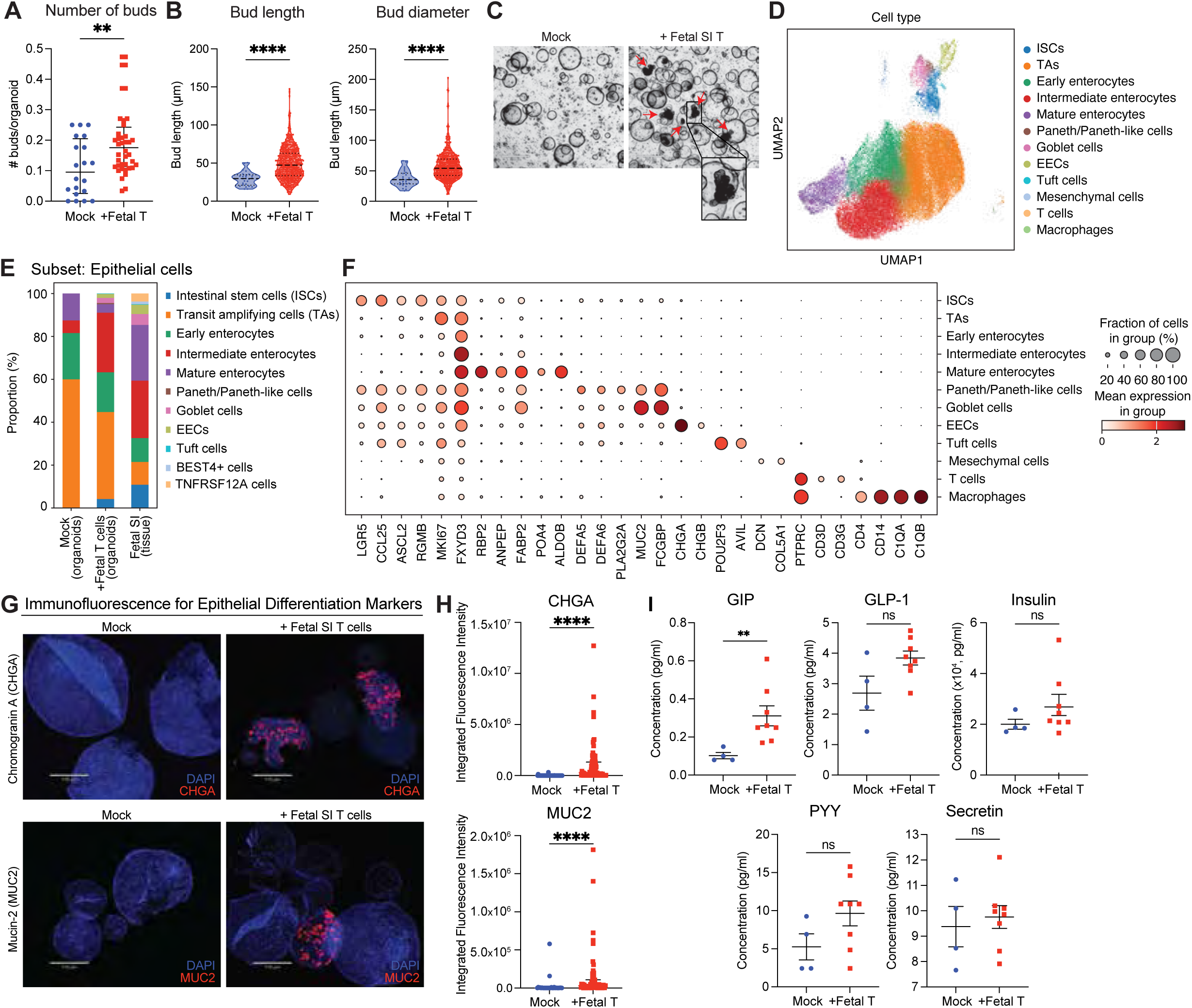
Fetal mucosal T cells promote intestinal stem cell commitment to the secretory lineage. Fetal SI organoids were co-cultured with fetal SI T cells for 6 days (2 days growth phase, 4 days differentiation phase). All analyses shown are from day 6. **(A-C)** Microscopy was performed on organoids, allowing measurement of bud number **(A)**, length, and diameter **(B)** as a proxy for organoid differentiation. **(C)** Representative microscopy images showing increase frequency of budding in organoids co-cultured with T cells. Arrowheads indicate examples of budded organoids. Scale bars are 530 µm. **(D-F)** scRNA-seq was also performed to identify cellular subtypes present in co-culture. **(D)** UMAP for identified cell clusters were then **(E)** subclustered on epithelial cells to identify relative cellular proportions in organoids grown with or without T cells, relative to physiologic cell frequencies in the fetal SI ^6^. **(F)** Dot plot guiding cluster annotations in **(D, E)**. **(G-H)** Organoids were stained by immunofluorescence (IF) for chromogranin A (CHGA) or mucin-2 (MUC2), indicative of enteroendocrine cells and goblet cells, respectively. **(G)** Representative images from IF staining and **(H)** quantification are shown. In **(H)**, each dot represents one field of view. Data are shown as medians ±IQR and were analyzed by Mann-Whitney. **(I)** Media were analyzed by Luminex to detect metabolic hormones associated with enteroendocrine cell functionality (GIP, GLP-1, insulin, PYY, and secretin). Data in **(A-B)** and **(I**, insulin) were analyzed by Mann-Whitney and are denoted as medians ±IQR. Data in **(I**, GIP, GLP-1, PYY, secretin) were analyzed via Welch’s t-test and are shown as means ±SEM. Significance is indicated as follows: * p<0.05; ** p<0.01; **** p<0.0001; ns, not significant (p>0.05). Experiments were performed with two T cell and organoid donors, and Luminex data were performed independently twice from two independent co-culture experiments with these same donors.

Unexpectedly, we identified the presence of mesenchymal and macrophage-like cells in these co-cultures by scRNA-seq (**Figure 2**; **Figures 3D and 3E**). We suspect that these cells are carry-over contaminants from T cell purification rather than organoid isolation, as they are present only in co-culture conditions. Upon integrating these clusters with scRNA-seq from fetal SI tissue, the macrophage-like cells did not map to any characterized myeloid population, though they are most like nonclassical intermediate (slan^-^) monocytes ^6^ (**Figures S4A and S4B**). This suggests that while ex vivo culture conditions maintained their viability, they may have phenotypically drifted from their original in vivo identity. Mesenchymal cells identified in organoid co-cultures, in contrast, mapped to true mesenchymal populations, as well as neural and endothelial populations (**Figure S4A, right**)^6^. To verify whether these cell types were major drivers of the observed organoid growth and differentiation, we performed organoid co-cultures with discrete cellular fractions: (1) T cell enriched fraction (CD3+; containing mostly T cells), (2) T cell depleted-myeloid enriched fraction (CD3-CD14+; containing mostly myeloid cells), or (3) T cell depleted-myeloid depleted fraction (CD3-CD14-; containing B cells/ILCs/mesenchymal cells/etc.) (**Figures S4C-H**). The T cell enriched condition resulted in significantly more organoids than either T cell depleted condition (**Figures S4C and S4D**). By bulk RNA-seq and pathway analysis, we observed an enrichment in differentiation programming only in the T cell containing fractions, highlighting the importance of T cells in mediating these processes (**Figures S4F and S4G**). Concordantly, given that the CD3 enriched fraction in these experiments still contained myeloid and mesenchymal contributors, we then assessed the sufficiency of T cells to independently drive ISC renewal and differentiation. We isolated CD3+ T cells from the fetal SI by fluorescence activated cell sorting and co-cultured them with fetal organoids (**Figures S4H and S4I**). Fetal organoids cultured with sorted fetal SI T cells still exhibited increased organoid generation, supporting that T cells can independently support ISC renewal. Overall, these data indicate that fetal T cells can orchestrate ISC renewal and commitment to the secretory lineage in the small intestine.

### Both fetal CD4+ and CD8+ SI T cells stimulate epithelial growth and differentiation

As we used positive T cell selection to isolate T cells for organoid-T cell co-cultures, they were seeded with T cells representing the relative frequencies found in the fetal SI tissue. To identify which T cell subtypes were present by the end of ex vivo cultures, we subclustered on T cells recovered from differentiation co-cultures and integrated them with the same sequencing dataset from the fetal SI (**Figures 4A and 4B**). Concordant with the fetal SI, more than 50% of the T cells from co-cultures were conventional CD4+ T cells (**Figure 4B**). Although both naïve and memory states were represented, CD4+ memory T cells were most populous, in agreement with other studies (**Figure 4B**)^6,10,12^. Additionally, we identified a small proportion of regulatory (Treg) and double negative (DN) T cells that were relatively comparable to their typical proportions in the fetal SI (**Figure 4B**). Cycling T cells were more prevalent in ex vivo cultures than in fetal tissue, though this may be a product of highly permissive culture conditions (**Figure 4B**). Interestingly, we saw a low frequency of CD8+ naïve T cells, and CD8+ memory cells were undetectable in co-cultures (**Figure 4B**). We did not recover other innate-like T cells, such as gamma delta T (γδT) or natural killer T (NKT) cells from co-cultures (**Figure 4B**). As CD4+ and CD8+ T cells represented the bulk of fetal T cells from the original co-culture inputs, we evaluated which of these conventional subsets underlie organoid generation and/or differentiation by directly sorting these cells from the fetal SI and co-culturing them with fetal organoids (**Figures 4C-F**). We seeded these co-cultures with CD4+ or CD8+ T cells at their relative proportions observed in the native fetal SI. While the magnitude was relatively weaker than with total T cells, surprisingly both CD4+ and CD8+ T cell subsets modestly enhanced organoid generation (**Figures 4C and 4D)**. These subsets trended toward weakly supporting enteroendocrine cell, but not goblet cell, differentiation in some organoids (**Figures 4E and 4F**). These data support that both conventional CD4+ and CD8+ T cells can direct ISC fate in the fetal intestine, though the magnitude of this effect is reduced relative to bulk T cells, where effects are potentially synergistic.

**Figure 4.**
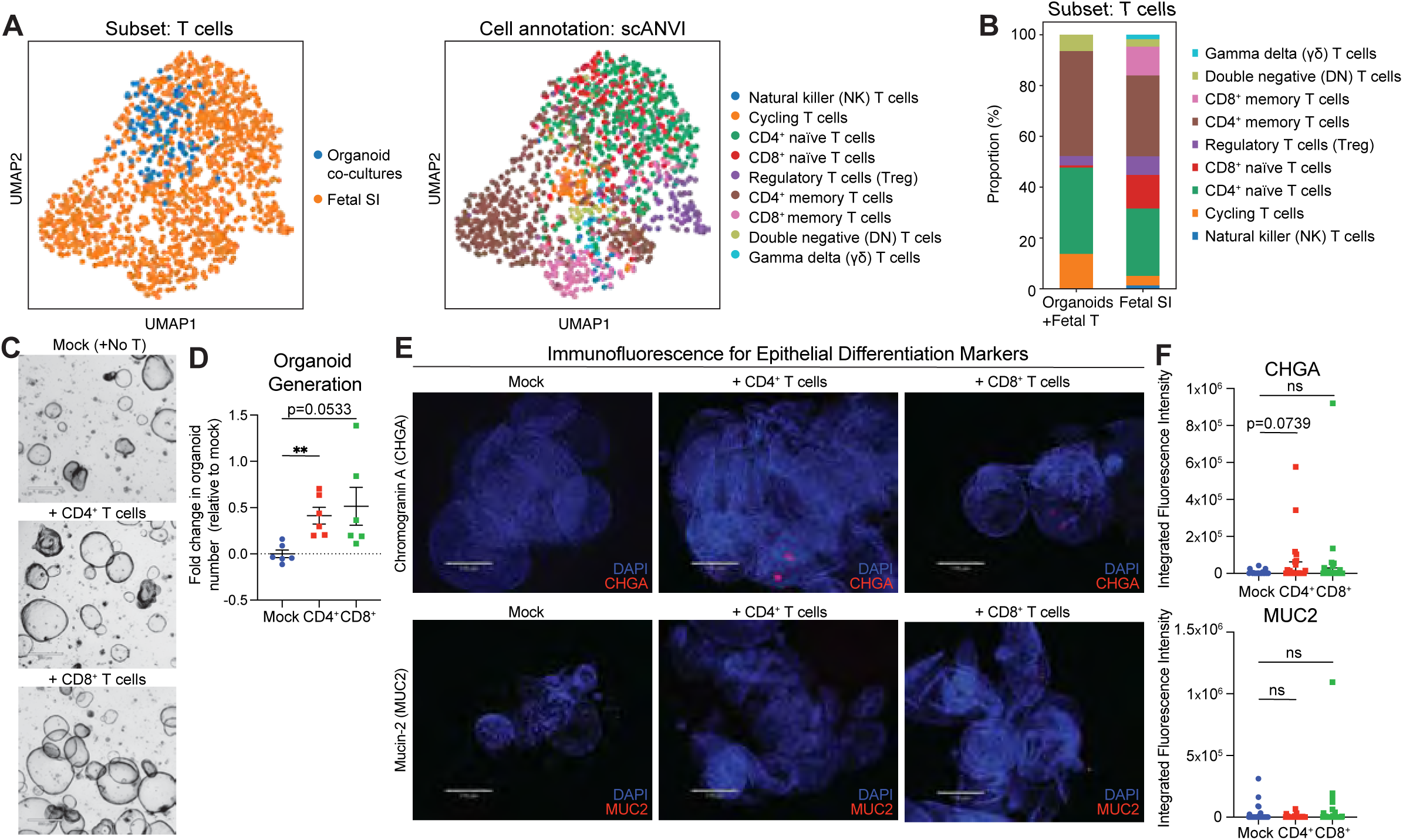
Fetal mucosal CD4+ and CD8+ T cells incite organoid generation and enteroendocrine cell differentiation. Single-cell RNA-sequencing data from differentiation phase organoid co-cultures were integrated with scRNA-seq from the fetal SI ^6^. **(A)** UMAP showing integration of T cells from organoid co-cultures or the fetal SI and annotated T cell subsets by single-cell annotation using variational inference (scANVI). **(B)** Proportion of each T cell subset identified in **(A)** from organoid co-cultures with T cells or the fetal SI. **(C, D)** T cells were positively selected, sorted into CD4+ and CD8+ subsets, and then co-cultured with fetal organoids. Independent T cell subsets were sufficient to induce organoid generation. Scale bars indicate 530 µm. Data in **(D)** are normalized to the number of organoids generated in the mock (no T cell) condition represented as fold change. Each dot represents one well of organoids. **(E-F)** Organoids were stained for chromogranin A (CHGA) or mucin-2 (MUC2), indicative of enteroendocrine cells and goblet cells, respectively. **(E)** Representative images from IF staining and **(F)** quantification are shown. Scale bars indicate 170 µm. In **(F)**, each dot represents one field of view. Data in **(D)** are shown as means ± SEM and were analyzed by Welch’s t-test. Data in **(F)** are shown as medians ±IQR and were analyzed by Mann-Whitney. Experiments in **(A-F)** were each performed once on different days with two independent fetal organoid and T cell donors. Significance is indicated as follows: * p<0.05; **p<0.01; ns, not significant.

### Secreted factors alone are not sufficient to drive organoid generation and differentiation

It has been described in adult mice that cytokines from CD4+ T cells can modulate ISC differentiation and self-renewal^26^. To determine whether cytokine(s) derived from human fetal T cells perform an analogous function, we assayed supernatant from fetal organoid cultures via multiplexed cytokine array for 96 human cytokines (**Figures 5A-D**). To maximize detection, we tested media at the first media change (day 3). Most cytokines enriched in co-cultures relative to mock were shared between fetal and adult SI T cell co-cultures, such as RANTES (CCL5), Granzyme A, CXCL9, sFasL, IL-2, and MIP-1β (CCL4), though some were only enriched in 2/3 adult T cell donors (**Figures 5A-D**). The presence of IL-2 in both fetal and adult co-cultures highlight the survival and functionality of both T cell age groups in organoid co-cultures (**Figures 5A and 5B**). Given that cytokine profiles were similar, we batch corrected these data to draw direct comparisons between fetal and adult T cells (**Figures 5C and 5D**). The cytokine profile of culture media clustered based on whether organoids were cultured without T cells, with fetal SI T cells, or with adult SI T cells (**Figure 5C**). In comparing fetal to adult SI T cell co-cultures, there were distinct differentially expressed cytokines (**Figure 5D**). Fetal T cell co-cultures were enriched for MIP-3β (CCL19), I-309 (CCL1), Granzyme B, MCP-2 (CCL8), MIP-1β, TARC (CCL17), and IL-33, whereas GM-CSF and IL-6 were enriched in adult T cell co-cultures (**Figure 5D**, left). Even with removal of the adult donor with lower overall cytokine detection, these same cytokines were enriched in fetal T cell conditions (**Figure 5D**, right).

**Figure 5.**
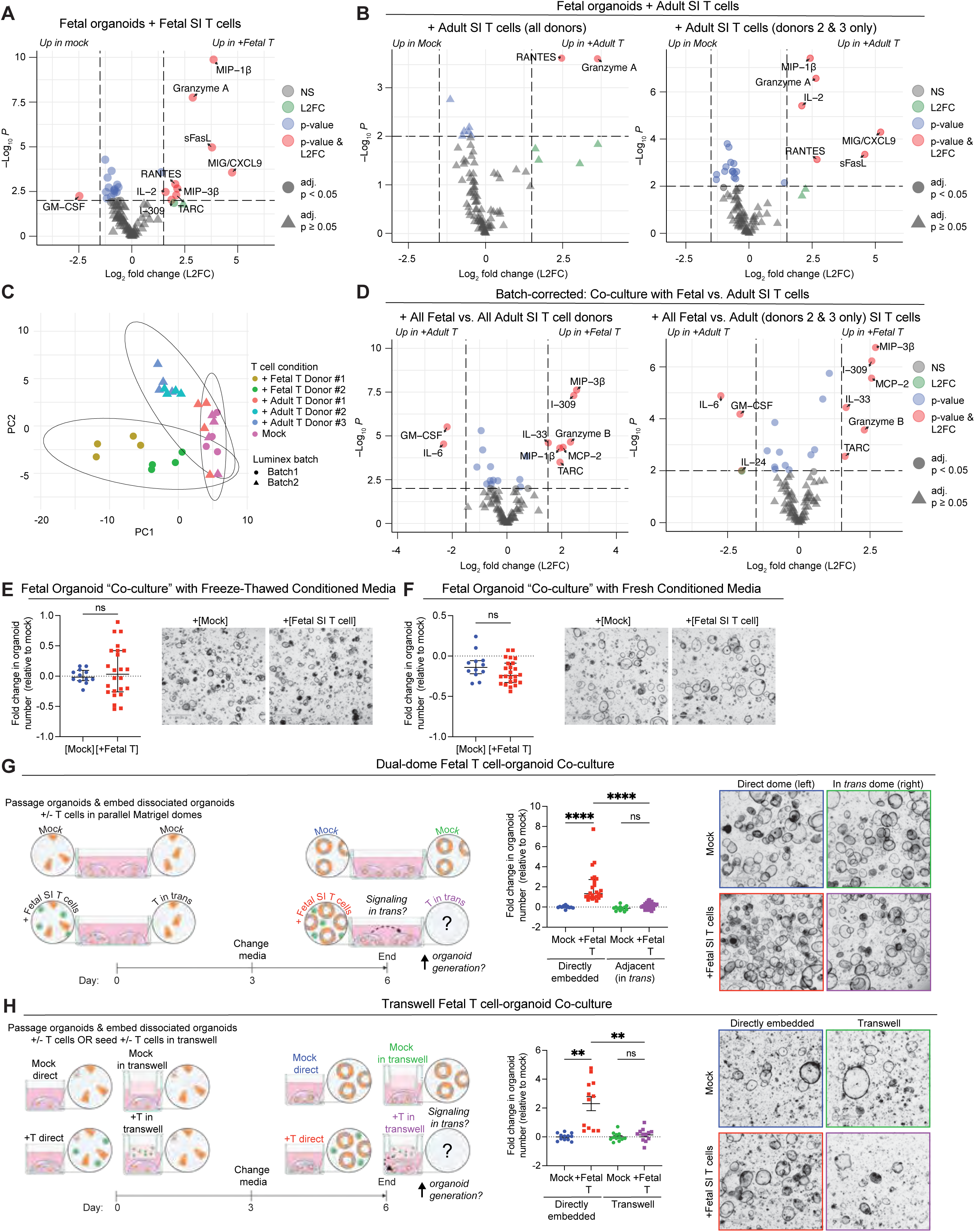
Secreted factors from fetal mucosal T cells are not sufficient to support organoid generation or differentiation. **(A-D)** Media from fetal SI organoids cultured with or without fetal **(A)** or adult **(B)** SI T cells were analyzed by Luminex 96-plex cytokine arrays on day 3 of culture. Volcano plots indicate enriched cytokines detected in co-culture with fetal SI T cells **(A)** or adult SI T cells **(B)** relative to mock conditions. Pairwise comparisons between all adult T cell donors indicate donor #1 clustered away from donors #2 and 3; therefore, volcano plots showing comparisons between mock and all adult T cell donors **(B, left)** or mock and only adult T cell donors 2 and 3 **(B, right)** are shown. **(C)** PCA for all Luminex data presented in **(A-B)** after batch-correction to account for inter-assay variability arising from independent multiplex cytokine array experiments. In **(A, B, D)**, |L2FC|>1.5 and p<0.01 were considered significant. Each experiment was independently repeated twice with these donors. **(E-F)** Fetal SI organoids were cultured in reconstituted conditioned media generated from fetal SI organoids cultured with or without fetal SI T cells. In **(E)**, conditioned media was collected on day 2, frozen, thawed and reconstituted before being added to newly passaged organoids. Images and organoid counts were collected on day 3. In **(F)**, fresh conditioned media were collected on day 2, concentrated, and reconstituted before being applied to newly passaged fetal SI organoids. On day 4, media from these conditioned cultures were collected again, concentrated, reconstituted, and used for a second round of treatment. Organoid images and counts were acquired on day 4. **(G-H)** To assess if fetal T cells can signal in *trans* via secreted factors, **(G)** fetal SI organoids were seeded in Matrigel^®^ domes adjacent to organoids embedded with or without T cells or **(H)** beneath transwells seeded with or without T cells. Organoid counts are normalized to the number of organoids generated in mock (no T cell) conditions, represented as fold change. Each dot represents one well of organoids. Images and quantification were collected on day 6. Experiments in **(E-H)** were independently performed twice using 1-2 fetal organoid donors and at least 2 fetal T cell donors. Scale bars in **(E-H)** are 530 µm. Data in **(E-F)** are shown as means ±SEM and were analyzed by Welch’s t-test **(E-F)** or one-way ANOVA with Tukey’s multiple comparisons test **(H)**. Data in **(G)** are shown as medians ±IQR and were analyzed via a Kruskal-Wallis with Dunn’s multiple comparisons test. Each data point represents one well of organoids. Statistical significance is indicated as follows: ** p<0.01; **** p<0.0001; ns, not significant.

To test whether these cytokines or other secreted factors could drive organoid generation and differentiation, we cultured fetal organoids with conditioned media from organoid cultures grown with or without fetal SI T cells (**Figures 5E and 5F**). To resupply necessary growth factors, we reconstituted concentrated supernatant from either mock or +T cell conditions with fresh media, then treated newly passaged organoids with it. Organoids cultured with thawed conditioned media from previous co-cultures did not increase generation (**Figure 5E**). However, cytokine bioactivity can be negatively impacted by freeze-thaw cycles. Therefore, we also treated organoids with fresh conditioned media, but this still failed to enhance organoid generation relative to “mock” conditioned media (**Figure 5F**). Given this unexpected result, we employed two additional orthogonal approaches to verify that secreted factors were not driving the proliferation phenotype (**Figures 5G and 5H**). First, we seeded dissociated organoids into two separate Matrigel^®^ domes within the same well, with only one dome containing embedded T cells (**Figure 5G**). Because small molecules and cells can move through Matrigel^®^, we reasoned that if secreted factor(s) from T cells promote stem cell renewal, then these factor(s) could potentially signal *in trans* to organoids in a distal Matrigel^®^ dome (**Figure 5G**, schematic). Second, to exclude competition between Matrigel^®^ domes as a potential sink for secreted signals, we cultured fetal organoids with fetal SI T cells suspended in a permeable transwell insert above them, allowing diffusion of soluble factors while precluding physical interactions (**Figure 5H**). In both conditions, organoid generation was significantly enhanced only if organoids were embedded directly with T cells, phenocopying the experiments with conditioned media (**Figures 5G and 5H**). Furthermore, we assessed whether T cells were able to promote organoid differentiation in these systems. Similar to organoid generation, we only observed the upregulation of genes associated with epithelial differentiation (e.g., *MUC5B*, *CHGA*) when T cells were directly embedded with organoids, but not when they were in transwells or adjacent Matrigel domes (**Figure S5**). Together, these data strongly suggest the necessity of physical interactions between fetal T cells and the epithelium to modulate intestinal stem cell fate.

### T cell proximity and signaling to the ISC niche decreases with developmental age

As fetal T cells do not exert these effects on ISCs by secreted factors alone, these data imply that they must either engage in direct cell contact and/or short-range paracrine interactions, which require close spatial proximity. We labeled T cells and measured their distance to the epithelium in co-culture with organoids (**Figures 6A and 6B**). While T cells localized near organoids in all conditions, by day 6, fetal T cells were more likely to be found near an organoid than adult T cells were (**Figures 6A and 6B**). We also observed cord blood T cells were the farthest away from organoids on day 2, though this distance decreases by day 6 (**Figure 6B**). However, growth-phase organoids lack crypt-villus architecture for identifying ISCs, and 2D projections of Z-stacks may not accurately capture spatial relationships. To verify these findings in vivo, we then evaluated the proximity of T cells to ISCs across the human lifespan by spatial transcriptomics profiling in fetal and neonatal (atresia/non-NEC and NEC) SI samples (**Figures 6C-I**).

**Figure 6.**
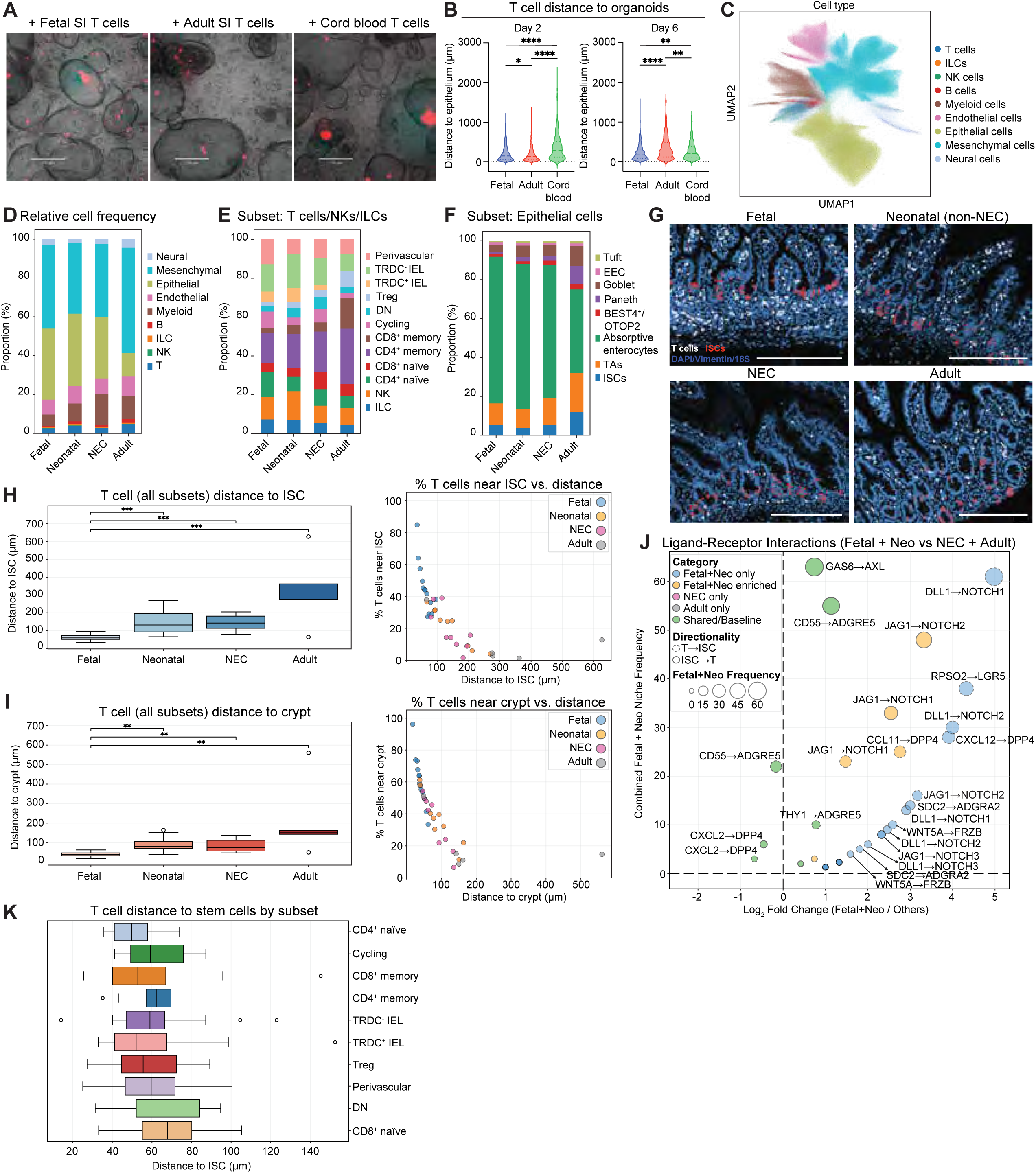
Fetal mucosal T cells localize closer to intestinal stem cells than adult or NEC-associated T cells. **(A-B)** T cells from the fetal or adult small intestine or cord blood were stained with CellTracker Red and co-cultured with fetal SI organoids. Representative images **(A)** and T cell distance to organoids **(B)** measured on days 2 and 6 of co-culture. Scale bars indicate 530 µm. **(C-J)** Single-cell spatial transcriptomics were conducted on fetal, neonatal (non-NEC), NEC, and adult SIs were performed. Cells were clustered by manual annotated guided by scRNA-seq of the SI ^6^, and distance between T cells and ISCs or cells in the crypt were tabulated. **(C)** UMAP of identified cell clusters from all age groups/conditions. **(D-F)** Relative proportions of cells occupying all clusters **(D)**, the T/ILC/NK cell cluster **(E)** and epithelial cell cluster **(F)** by donor age group/condition. **(G)** Representative images of T cell and ISC localization in Xenium Explorer, where scale bars indicate 200 µm, from the spatial transcriptomics dataset. **(H)** Distance from T cells to ISCs (left) and % T cells near ISC as a function of distance (right). **(I)** Distance from T cells to crypts (left) and % T cells near crypts as a function of distance (right). **(J)** Ligand-receptor interactions predicted by CellChat between T cells and ISCs, shown as interactions enriched in fetal and atresia neonatal donors relative to adult or NEC donors. Niche frequency refers to the number of crypts the interaction was identified in. Interactions found in only one niche were excluded. **(K)** T cell distance to ISCs separated by T cell subset. Data in **(B)** are presented as medians and upper and lower quartiles and were compared by Kruskal-Wallis with Dunn’s multiple comparisons test. Data in **(H-I**) are presented as medians ± 1.5 IQR and were analyzed by t-test **(H)** or Mann-Whitney **(I)** with Bonferroni correction. Significance is indicated as follows: * p<0.05, ** p<0.01, *** p<0.001. No significance annotation indicates statistically insignificant comparisons relative to fetal donors.

Using previous scRNA-seq^6^ data to guide cell clustering, we identified a diverse cellular landscape of immune, endothelial, epithelial, mesenchymal, and neural cells in these SI sections (**Figures 6C-F; Figure S6**). We identified comparable frequencies of secretory cells, including a postnatal increase in goblet and Paneth cells (**Figure 6F**)^6^. By spatial transcriptomics, we detected a larger proportion of mesenchymal cells and fewer T cells, possibly due to differences in cell isolation (e.g., dissociation versus sectioning) between these methods (**Figure 6D**). Nonetheless, this dataset supports previous studies on the developmental timeline of T cell emergence in the SI, with conventional T cells being detectable as early as 12 weeks EGA (**Figures S6E and S6F**). We similarly identified an increase in both T cell abundance and subset diversity throughout the second trimester, particularly CD4+ T cells while CD8+ memory T cells expanded postnatally (**Figures S6E and S6F**). In agreement with our prior work, early-life T cells were enriched for transcripts associated with stem-like signatures, including *ZBTB16* (encoding PLZF) and *IKZF2* (encoding Helios) (**Figure S7A**). In addition to conventional T cells, we found innate-like and conventional T cells with memory states and frequencies concordant with our previous reports^6,10^ (**Figure 6E**).

To examine how T cells and epithelial cells may interact in the bona fide intestinal epithelium, we examined T cell distance relative to the stem cell niche. Overall, we observed an abundance of intraepithelial and submucosal T cells in fetal donors relative to postnatal donors (**Figure 6G**). When quantified across all donors, fetal T cells were significantly closer to ISCs than were T cells from neonates or adults (**Figures 6H and 6I**). ISCs are often reported to be more prevalent in fetal tissue^27^, but sections used for these spatial transcriptomics experiments indicated more ISCs in adult donors (**Figure 6F**). Fetal T cells were enriched around ISCs and crypts relative to neonatal, NEC-associated, and adult T cells (**Figures 6H and 6I; Figures S7B and S7C**). We identified similar trends for CD4+ and CD8+ T cell subsets, though there were few CD8+ memory T cells in prenatal tissue (**Figures S6F and S6G; Figures S7B and S7C**). Strikingly, NEC T cells less densely occupied the stem cell niche despite being close in developmental age to the fetal and neonatal donors (**Figures 6H and 6I; Figures S7B and S7C**). This reduction in T cell density near the crypt occurred even though NEC is characterized by substantial villus blunting, a feature that would be expected to compress tissue architecture and potentially increase T cell-ISC proximity^13^. Overall, CD4+ naïve and CD8+ memory T cells were positioned nearest to ISCs (**Figure 6K**). These findings suggest that T cell positioning within the ISC niche is not a passive consequence of intestinal architecture, but rather an actively regulated developmental program that is disrupted in disease.

We next segmented tissue sections into functional niches to focus on cell-cell communication within intestinal crypts, where ISC maintenance is coordinated. We identified a strong enrichment of ligand-receptor pairs involved in Notch and Wnt signaling in early-life donors, with interactions including *DLL1-NOTCH1/2/3*, *JAG1-NOTCH1/2/3*, *RSPO2-LGR5*, *WNT5A-FRZB,* and *SDC2-ADGRA2* (**Figure 6J**). These interactions were largely absent in adult and NEC sections (**Figure 6K**), indicating a developmentally restricted signaling network within the crypt niche. Many of these factors were expressed on both T cells and ISCs with predicted signaling interactions in both directions, consistent with bidirectional crosstalk between these cell types (**Figure 6K**). Given the established roles of Notch and Wnt pathways in regulating ISC self-renewal and differentiation, these findings suggest that early-life T cells participate directly in shaping the regenerative crypt environment. Together, these data demonstrate that healthy early-life T cells are developmentally programmed to occupy the ISC niche where they support appropriate intestinal growth and differentiation, whereas this is disrupted in diseased states, such as NEC.

### Fetal SI T cells can rescue proliferation and differentiation in the NEC epithelium

To determine if these T cell-ISC interactions are implicated in disease pathology, we next assessed the functional capacity of NEC T cells to support epithelial growth and differentiation. In addition to disrupted T cell localization, NEC is associated with inflammatory T cell signatures and reduced goblet and enterocyte populations^13,28–30^. Inflammatory CD4+ T cells from the intestine in NEC mouse models can induce tissue damage via inflammatory cytokine production, but how T cells contribute to NEC in the human intestine is poorly defined^31^. This led us to ask whether NEC ISCs are intrinsically defective at supporting epithelial growth and differentiation, or whether T cells mediate damage either directly (e.g., impeding these functions) or indirectly (e.g., failing to provide appropriate supportive cues. First, we compared ISCs from fetal or neonatal (non-NEC) donors versus with those with NEC (**Figures 7A and 7B**). Relative to NEC-associated ISCs, fetal or neonatal ISCs were enriched for stem cell identity (*ASCL2*, *CD44*, *GATA6*), proliferative programming (*MCM3/4/5*), regeneration (*YAP1)*, growth factor signaling (*FGFR4*, *ERBB2/3, PIK3R3*), Wnt and cell fate signaling (*AXIN1, SMAD4*), structural integrity (*EPCAM*, *CDH1*) (**Figures 7A and 7B**). *CCL25*, a chemokine involved in immune cell recruitment that we also identified by bulk RNA-seq (**Figures 2A and 2B**), was significantly enriched in fetal and neonatal ISCs relative to NEC-associated ISCs (**Figures 7A and 7B**). To compensate for NEC-associated T cells being found farther away from ISCs in vivo, we used our in vitro organoid co-culture system to position NEC T cells near ISCs. Therefore, we co-cultured organoids from fetal or NEC donors with T cells from these same donors. T cells from the SI of an infant with NEC failed to augment organoid generation in both the NEC-derived and fetal SI organoids, suggesting that these NEC-associated T cells additionally lack the factors needed to support organoid generation. In line with this model, fetal SI T cells co-cultured with NEC-derived organoids could rescue this defect at a magnitude comparable to their effect on fetal organoids (**Figures 7C and 7D**). Moreover, NEC T cells exhibited diminished capacity to induce enteroendocrine cell differentiation relative to fetal T cells (**Figures 7E-H**) and trended this same direction for goblet cell differentiation, though this was not statistically significant (**Figures 7F-H**). In addition to restoring proliferation, fetal SI T cells also induced secretory lineage differentiation in organoids from fetal and NEC donors (**Figures 7E-H**). Taken together, these data suggest that NEC T cells may contribute to disease by failing to support ISC renewal and differentiation, implicating disrupted T cell-epithelial crosstalk as a contributor to NEC pathogenesis. These data importantly suggest that NEC-associated tissue can be driven towards proliferation and secretory lineage differentiation when appropriate signals are provided.

**Figure 7.**
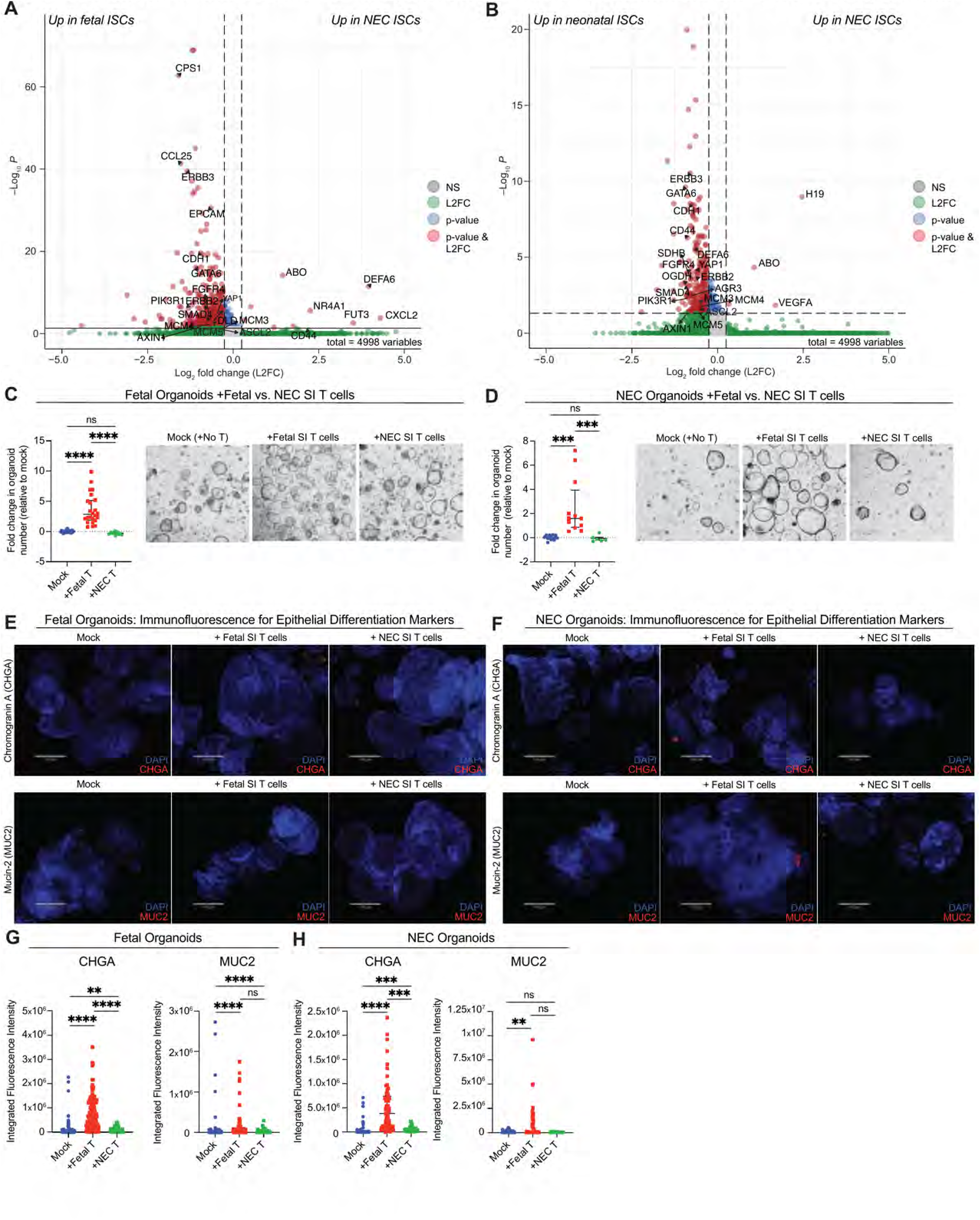
Fetal mucosal T cells can rescue proliferation and differentiation defects in the NEC epithelium. **(A-B)** Volcano plots generated from spatial transcriptomics data showing differentially expressed transcripts ((|L2FC|>0.5, p_adj_<0.05) in ISCs in fetal **(A)** and neonatal (non-NEC) **(B)** donors relative to NEC donors. **(C-H)** Fetal or NEC organoids were co-cultured with fetal or NEC SI T cells. **(C-D)** Fetal organoids **(C)** and NEC organoids **(D)** were counted and imaged on day 6 of culture. Organoid number is shown as fold change relative to mock conditions, where each dot indicates one well of organoids. Scale bars are 530 µm. **(E-H)** Organoids were stained for chromogranin A (CHGA) or mucin-2 (MUC2), indicative of enteroendocrine cells and goblet cells, respectively. **(E-F)** Representative images from IF staining from fetal organoids **(E)** and NEC organoids **(F)**. Scale bars indicate 170 µm. **(G-H)** Quantification for CHGA and MUC2 staining. In **(G-H)**, each dot represents one field of view. Data in **(C-D, G-H)** are shown as medians ±IQR and were analyzed by Kruskal-Wallis with Dunn’s multiple comparisons test. Each experiment was independently performed twice using 1-4 fetal SI T cell donors, 1 NEC T cell donor, 2 fetal organoid donors, and 1 NEC organoid donor. Statistics are indicated as follows: ** p<0.01; *** p<0.001; **** p<0.0001; ns, not significant.

## DISCUSSION

Adaptive immune cells and their dynamic shifts through gestation and postnatal development have only recently been characterized in the human intestine. Despite their abundance and diversity, few studies have linked these cells to functional roles in this mucosal niche, even though lifelong mucosal function is established beginning with epithelial development in utero. This gap offers an invaluable opportunity to define how immune cells actively shape, rather than simply respond to, epithelial development. Using a T cell-organoid co-culture model derived from fetal, neonatal, and adult small intestinal tissue, we identified that fetal and neonatal mucosal T cells can modulate intestinal stem cell (ISC) fate, even in the adult epithelium. This demonstrates a previously unrecognized capacity of early-life T cells to exert instructive, developmentally coordinated effects on the epithelium. Specifically, T cells from healthy early-life (e.g., fetal and neonatal) small intestinal tissue support ISC renewal and differentiation. In contrast, T cells derived from cord blood were insufficient to activate these pathways, indicating that these effects are more context-specific rather than a general property of all early-life T cells. Even in growth phase, T cells directed differentiation along the secretory lineage, suggesting that T cells supply critical differentiation factors that are absent in traditional growth media.

Differentiation was further augmented when cultures were maintained in differentiation media, suggesting that additional components (e.g., Notch inhibitors, reduced Wnt) further facilitate this process. ISC renewal and differentiation are classically viewed as opposing, tightly regulated processes. However, the epithelium can also achieve differentiation through “regress to progress” strategies that rely on augmenting ISC number and potency^23^. We propose that healthy early-life T cells activate a similar program by activating proliferative and anti-apoptotic programming in ISCs, ultimately increasing total stem cell numbers and enabling downstream differentiation. This model provides a unifying framework linking immune signals to coordinated expansion and maturation of the epithelial compartment. Our data further support that, rather than driving epithelial damage, NEC-associated T cells are missing critical interactions with ISCs, at least in part because of their distance from the crypt niche. More broadly, our study establishes early-life mucosal T cells as key determinants of epithelial fate and function, highlighting T cells and their interactions with ISCs as potential targets for regenerative and immunomodulatory therapies. Specifically, these applications span not only NEC, but also inflammatory bowel disease and short bowel syndrome, where epithelial renewal and/or differentiation are also largely dysregulated.

While our data support that early-life T cells can independently drive these processes, we cannot rule out the possibility that mesenchymal and myeloid cells cooperate with T cells in co-culture or *in vivo*. Fetal mesenchymal cells adjacent to proliferative crypts produce neuregulin 1 (NRG1), and NRG1 can support epithelial growth and cellular diversification in freshly isolated crypts^5^. However, NRG1 cannot drive renewal and differentiation in long-term expanded organoids^5^, and all organoids used in our study were maintained for at least three passages before co-culture with CD3+ fractions. There is active crosstalk between T cells, myeloid cells, and mesenchymal cells with each other and with the epithelium in neonates, and these interactions are disrupted in NEC^13^. Indeed, mesenchymal stromal cells can support epithelial regeneration, and impairment of this stromal-epithelial crosstalk is associated with Hirschsprung-associated enterocolitis^32^. Given that T cells weakly supported enteroendocrine differentiation and failed to promote goblet cell emergence, it is likely that these cells engage in multicellular crosstalk to support the ISC niche. It is also possible that goblet cell identity requires additional factors present only in differentiation media, since goblet cell differentiation was most robust in differentiation phase. This is also consistent with our spatial findings regarding NEC-associated ISCs, as these cells retained core stem cell identity while displaying altered epithelial-immune signaling signatures. These data collectively support that other epithelial, stromal, or innate immune compartments also provide signals in healthy development and in NEC.

Though our data reveal that T cells can support organoid generation and differentiation, it is not entirely clear from a mechanistic standpoint how they do this or why NEC and adult T cells fail to perform these functions. One possibility is that T cells in these states are localized too far away from the ISC compartment to provide these cues, which is supported by our spatial transcriptomics data. However, T cells may also prioritize more classical immune surveillance functions in adulthood, characterized by inflammatory profiles, whereas the intestine before or during early postnatal microbial exposures has other developmental demands. In utero, the intestine must dramatically grow and differentiate to prepare for early microbial exposures. Between 19-23 weeks EGA – when T cells most dramatically expand in the fetal SI – the SI lengthens by roughly 34%, with growth decelerating in infancy and after the first year of life^33,34^. However, T cell populations between early-life and adulthood are distinct, which could explain the differences observed between early-life and older T cells in our co-cultures. For example, the early-life SI harbors substantially more naïve T cells, though CD4+ memory T cells are the most abundant by the mid-second trimester^6,10,12^. Fetal and neonatal SI T cells exhibit striking similarities in both abundance and expression profiles. The adult SI, in contrast, is most heavily populated by CD8+ memory T cells, which are present but scarce early in development. A similar possibility should be considered for NEC T cells, which exist in different proportions than is typical of the healthy developing SI. Specifically, naïve T cells are enriched in NEC, while memory T cells are decreased^6^. NEC-associated naïve T cells are transcriptionally similar to those from fetal tissue, but NEC-associated memory T cells are much more transcriptionally concordant with later developmental ages (e.g., pediatric and adult)^6^. Notably, human fetal CD4+ memory T cells have been previously implicated in supporting ISC proliferation, but not differentiation, via production of tumor necrosis factor alpha (TNFα)^10^. Therefore, it is conceivable that T cells in adults and in NEC simply exist in the wrong proportions in co-cultures to carry out these functions. Nonetheless, fetal and neonatal T cells do exhibit unique features. Our spatial data support previous reports of early-life T cells being highly enriched for stem-like signatures^6,35,36^. Similarly, memory T cells in NEC and adults are more enriched for activation-associated genes, including *GZMB*^6^.

Despite higher proportions of memory CD8+ cells and cytotoxicity programming in adult T cells, we interestingly observed elevated levels of granzyme B in organoids co-cultured with fetal T cells relative to those grown with adult T cells. Unexpectedly, we did not detect differential expression of TNFα between T cell and mock conditions by Luminex, suggesting an alternative mechanism at play. One explanation for this disparity is that TNFα-mediated ISC proliferation was identified in donors with a maximal EGA of 20 weeks, preceding abundant T cell expansion in the fetal SI^12^. Studies in adult mice have demonstrated that T helper cytokines from Th1, Th2, Th17, and Treg cells can support ISC differentiation and self-renewal^26^. IL-22 derived from neonatal Th17 cells has also been shown to support organoid generation and differentiation^19^. However, we did not identify these cytokines in our T cell populations or co-cultures either. Moreover, the fetal intestine is strongly biased toward Th1-like T cells, which failed to promote secretory fates or stem cell renewal at least in mice^6,10,26^.

Our data strongly also suggest that T cells must physically interact with, or reside in very close proximity, to ISCs to promote growth and differentiation. T cell trafficking patterns and predicted ligand-receptor interactions further support this notion. In mapping T cell positioning relative to ISCs, we observed that fetal T cells were significantly closer to stem cells when compared to other age groups, suggesting preferential localization within the ISC niche during early life. Fetal T cells also expressed distinct ligand-receptor pairs with cognate partners detectable on ISCs, suggesting direct communication with the stem cell compartment and potential for direct signaling interactions that may modulate cell behavior. However, in NEC, T cells were situated farther from crypts, impeding T cells from providing these developmental cues to ISCs. Therefore, our data highlights that NEC-associate T cells not only exhibit distinct expression profiles from healthy early-life T cells, but they also fail to appropriate localize to the stem cell niche. These interactions reveal developmentally restricted immunomodulation of ISC fate by activating Wnt and Notch signaling. *DLL1*, *JAG1*, *NOTCH1/2,* and *SDC2* transcripts have been independently detected in fetal intestinal T cells by single-cell sequencing^27^ and Notch ligand expression has been reported on activated T cells^37,38^. Although fetal intestinal T cells are not all activated, many are effector memory skewed and show evidence of clonal expansion, which may explain baseline expression of these markers in at least some cells^10,12,39^. It is possible that some of these interactions are proximity artifacts rather than being actively expressed in T cells, such as *WNT5A* and *RSPO2*, as these are more classically associated with stromal and epithelial sources.

We cannot exclude that physical interactions induce soluble cues from T cells to mediate these processes. In mice, T cell modulation of ISC fate additionally required cell-cell contact between the T cell receptor (TCR) and major histocompatibility class II (MHCII)^26^. MHCII is expressed on human ISCs *in vivo* and in organoid models^40^, but our co-cultures did not require donor-matching, enabling the age-mismatching experiments presented here. It is conceivable that TCR could associate with MHCII based on conserved structural properties rather than haplotype-dependent MHC:peptide interactions. MHC molecules were also not predicted interactions in our spatial dataset, implying an alternative method of T cell-ISC communication. Thus, our data point to an alternative mechanism for T cell modulation of ISC fate during early human development, warranting future investigation. Regardless, our study establishes a donor-matched and -mismatched human co-culture system for interrogating MHC-independent T cell-ISC interactions, providing a scalable framework to dissect age- and tissue-specific crosstalk.

Finally, our study is not without limitations. Tissue from infants can only be ethically collected when there is medical concern, meaning that the infant tissue included in this study is derived from diseased patients. To minimize this inherent limitation, we utilized infants diagnosed with intestinal atresia, where disease etiology is not classically immune or epithelial in origin, except for NEC cases^41^. In atresia and adult diseased cases, we used adjacent healthy tissue collected during resection wherever possible. For NEC, most cases occur in infants born before 32 weeks EGA, and its incidence is higher and more severe in extremely preterm infants (<28 weeks EGA). For this reason, fetal donors are the closest age-matched healthy controls for comparison, but they lack bona fide microbial colonization typical of postnatal life. Collection of fetal tissue without congenital anomalies after 23 weeks EGA is unethical. Therefore, we acknowledge that these comparisons cannot be perfectly age- or disease-matched. To generalize our findings despite these limitations, we performed organoid co-cultures with multiple organoid and T cell donors, including these donors for spatial transcriptomics when embedded sections existed, which all showed similar trends. However, we cannot exclude the possibility that there is donor-specific variation, or that differences in abundance and proportions of cells were secondary biases from tissue collection and sectioning. Although our annotation strategy successfully resolved distinct T cell subsets, we observed a subset of cells displaying ’chimeric’ expression profiles – such as the detection of EPCAM in intraepithelial lymphocytes (IELs) or mesenchymal markers in perivascular T cells. This is in part due to the known constraints of deep learning-based segmentation in complex tissues, where the significantly smaller size of T cells relative to adjacent epithelial or mesenchymal cells complicates precise boundary delineation. By visualizing the physical spatial distribution of these T cells within Xenium Explorer, we confirmed that their localization – strictly within the crypt/villus or submucosal compartments – aligns with their identified phenotypes, corroborating our findings despite the inherent limitations of segmentation models.

Ultimately, our study highlights T cells as active drivers of coordinated epithelial programming in the human intestine, whose interactions with the epithelium shift across distinct developmental life stages. Early-life T cells uniquely shape ISC fate decisions, underscoring that immune-epithelial crosstalk plays a formative role in establishing and maintaining epithelial homeostasis. While a similar process is mediated by myeloid cells in the lung^9^, our study reveals a nonclassical role for T cells as active shapers of intestinal development. In disease contexts such as NEC, epithelial integrity is compromised during a critical developmental window, and our study indicates that NEC-associated T cells fail to appropriately instruct ISC behavior. Impaired gut motility is not just as a symptom of NEC^42^, but it is also likely a critical factor in its development. Although peristalsis is decreased in preterm infants overall, our study suggests that this is potentially secondary to altered intestinal T cell dynamics, leading to decreased differentiation of secretory cells and their associated factors that support food digestion and motility. We propose that decreased motility can then create a permissive environment for bacterial overgrowth leading to downstream inflammatory cascade associated with NEC^13,28–30^.

These findings highlight the utility of strategies targeted at restoring regeneration and differentiation rather than solely mitigating inflammation, therefore offering broad insights to approaching therapeutics for intestinal disease. More broadly, our finding that adult ISCs remain responsive to early-life developmental immune cues reveals an unexpected degree of epithelial plasticity well into adulthood. This raises the possibility that developmental programs normally restricted to early life could be therapeutically reactivated to enhance epithelial repair and regeneration following disease, injury, or resection. This is particularly compelling for short bowel syndrome, a malabsorptive state caused by extensive intestinal loss, most commonly following surgical resection from NEC in infants or inflammatory bowel disease in adults. Defining the conditions under which these programs can be effectively engaged without uncontrolled cellular proliferation is an exciting avenue for future investigation, with major implications spanning both intestinal diseases and regenerative medicine.

## METHODS

### Tissue sample collection and cryopreservation

Human small intestinal tissue and cord blood samples were collected at Yale School of Medicine with informed consent and IRB approval (IRB #2000028360/2000028415; HIC #2000025173) or at the University of Pittsburgh. For samples collecting at Pittsburgh, tissue was collected under a discarded specimen protocol under IRB approval (IRB# PRO17070226). Cord blood was processed with two sequential red blood cell lysis steps using 1X RBC lysis buffer (Invitrogen, cat#00-4333-57) according to manufacturer’s protocol before cryopreservation. Samples were cryopreserved in 10% DMSO + 90% heat inactivated Fetal Bovine Serum (HI FBS) (Corning). All donors are outlined in **Table 1**. H291, H301, and H341 adult organoid lines were obtained from the Harvard Stem Cell Core under Material Transfer Agreement.

**Table 1.**
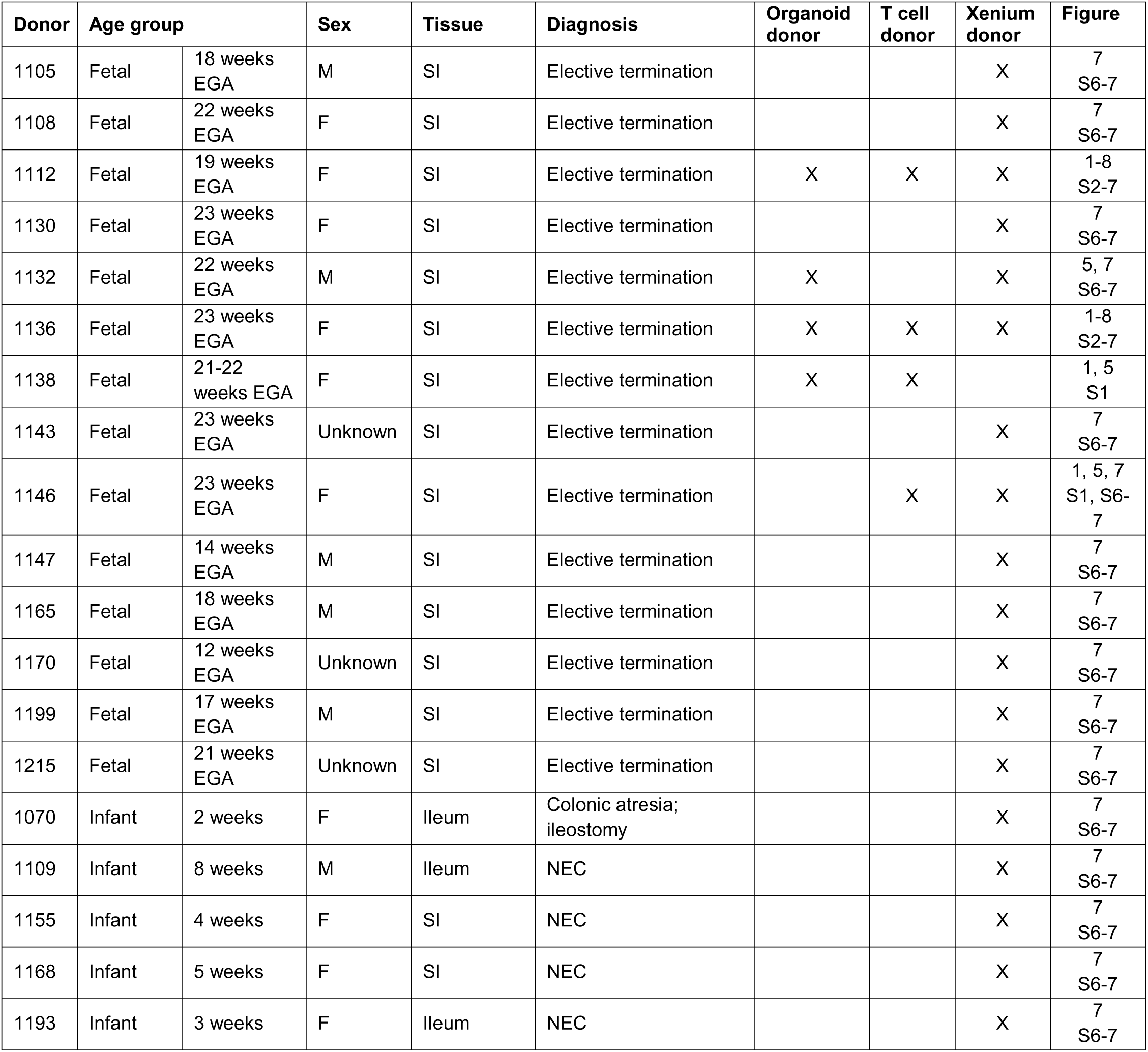
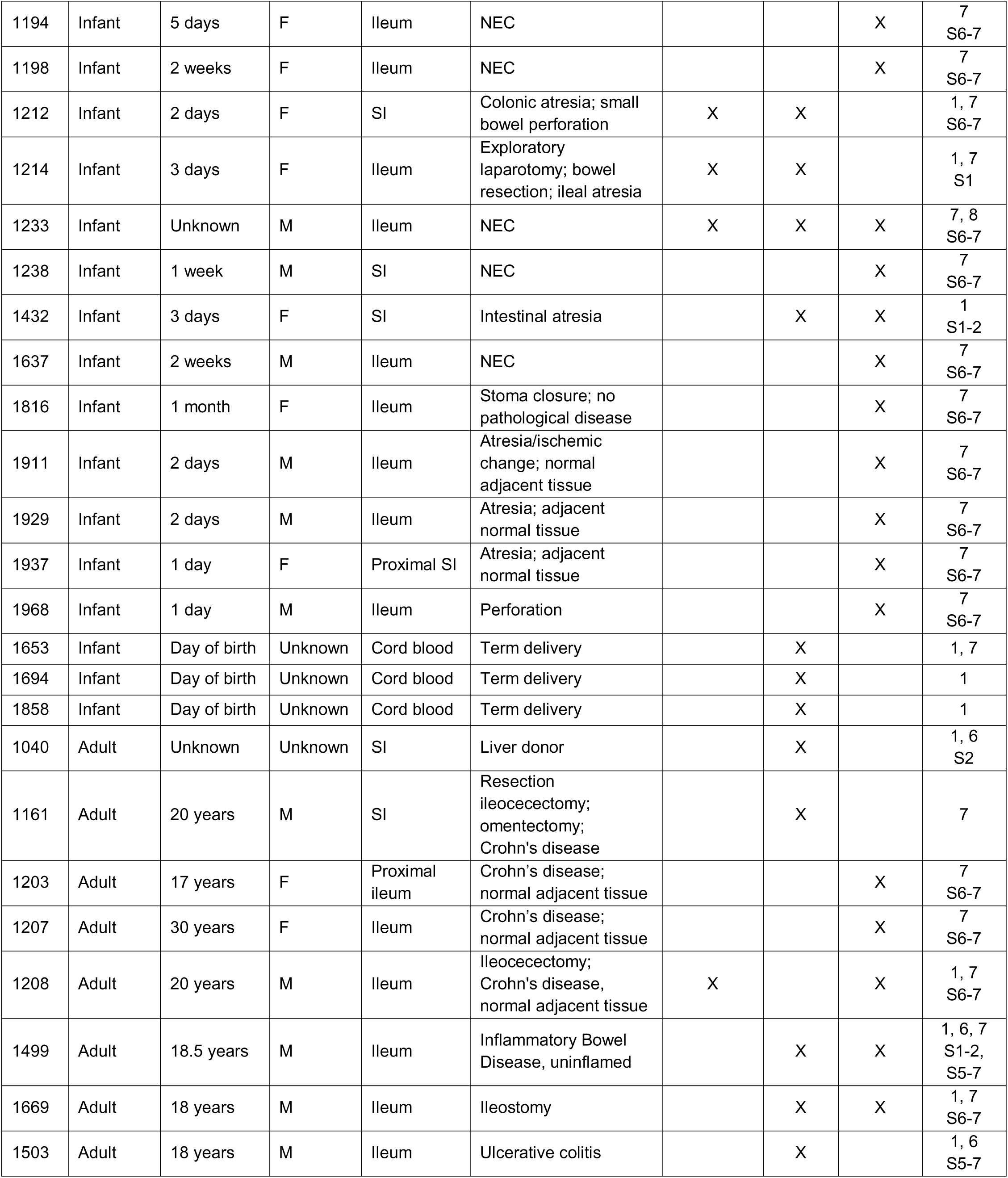
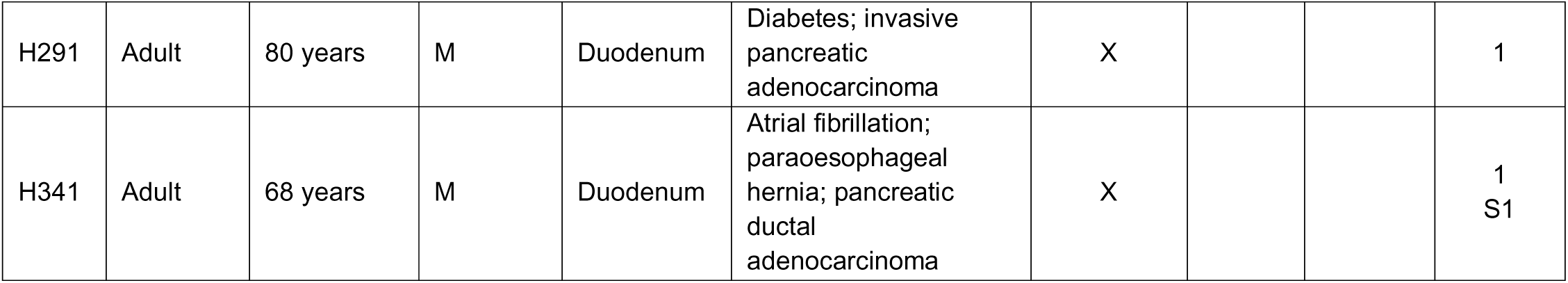
Donor metadata for all organoid and T cell donors utilized in the study.

### Organoid isolation

Organoids were generated by crypt isolation from cryopreserved small intestine tissue by StemCell Technologies manufacturer’s protocol, adapted from early intestinal organoid models derived from single LGR5+ stem cells^1^. Briefly, cryopreserved tissues were washed once each with 1X PBS (Gibco) and Advanced DMEM/F12 (Gibco, cat#12634010). Crypts were dissociated from tissue using Gentle Cell Dissociation Reagent (StemCell Technologies, cat#100-0485) on ice for 30 mins, followed by straining over 70 µm cell strainers. Crypts were pelleted at 290xg/5 min/4°C before plating in 25 µl Growth-Factor Reduced Phenol-Free Matrigel^®^ (Corning cat#356231) in 48-well plates (CytoOne, cat#CC7682-7548). After basement membrane polymerization, embedded crypts were given 250 µl IntestiCult Human Organoid Growth Media (StemCell Technologies, cat#06010) supplemented with 1% Penicillin/Streptomycin (Penn/Strep) (Gibco), 1% Amphotericin B (Gibco), and 10 µM Y-27632 (StemCell Technologies, cat#72304) per well. Initial organoid formation and growth proceeded for 10-20 days with media changes every other day before initial passage. Organoids were maintained at temperature- and humidity-controlled conditions of 37°C/5% CO_2_ and were passaged at least three times before use in experiments.

### Organoid maintenance and passaging

Organoids maintained in 3D were passaged every 5-8 days according to manufacturer’s protocol. In brief, organoids were collected in Gentle Cell Dissociation Reagent and incubated for 10 minutes at room temperature, with gentle rocking (∼60-70 rpm). Organoids were pelleted at 290xg/5 min/4°C and subsequently mechanically dissociated with vigorous pipetting in cold Advanced DMEM/F12. Dissociated cells were strained over a 70 µm filter and centrifuged at 500xg/5 min/4°C. Organoids were plated in Matrigel^®^ for a final passage ratio of approximately a 1.5 – 2-fold passage. Each well was given IntestiCult Human Organoid Growth Media containing Y-27632 and Penn/Strep in the above ratios (henceforth referred to as OGM), with media replacement every other day. Once per week, cultures were confirmed to be negative for mycoplasma using MycoStrip (InvivoGen, cat#rep-mys-10). Organoids were cryopreserved using freezing media consisting of 10% DMSO/70% OGM/20% HI FBS, per manufacturer’s recommendation. After thaw, organoids were passaged at least three times before use in experiments.

### T cell-organoid co-cultures

#### T cell isolation

Cryopreserved intestinal tissue was washed twice sequentially. First, tissue was washed with T cell media (i.e., RPMI (Gibco) containing 10 mM HEPES (Gibco), 1 mM sodium pyruvate (Gibco), 1% MEM Non-Essential Amino Acids (Gibco), 1% GlutaMAX (Gibco), 1% Penn/Strep, and 10% HI FBS). Second, tissue was washed with 1X PBS. Minced tissue was digested overnight in 25 mL T cell media containing 1 µg/mL DNase (Sigma Aldrich, cat#11284932001) and 8 µg/mL collagenase IV (Sigma Aldrich, cat#C5138-100MG) for overnight for approximately 16 hours with heated agitation (300 rpm/37°C) in MACS C tubes (Miltenyi). The next morning, tissue was digested for 1 hour using the OctoMACS 37°C heated and automated digest (Miltenyi). To remove particulate debris, digests were strained over a 70 µm cell strainer and pelleted at 300xg/10 min/4°C before cell counting. T cells were isolated via CD3 microbead positive selection (Miltenyi, cat#130-050-101 or 130-097-043) via MS or LS columns (Miltenyi, cat#130-042-201 or 130-042-401) per manufacturer’s instructions (Miltenyi). Once isolated, cells were resuspended in T cell media, were counted, and were stored at 37°C/5% CO_2_ during organoid passaging. Organoids were passaged as described above and combined with 5,000 positively selected T cells per well, unless otherwise stated. Each experiment contained organoids plated with or without T cells in three technical triplicate wells. Organoids were cultured for 6 days in IntestiCult OGM lacking Y-27632 (referred to as OGM(-Y)), unless otherwise stated, with one media replacement on day 3. All co-cultures were grown and maintained at 37°C/5% CO_2_.

#### T cell-organoid co-culture live imaging

Prior to seeding T cells in co-culture, apoptotic cells were removed via dead cell selection (Miltenyi, cat#130-090-101), and T cells were enriched via CD3-positive selection as described above. CD3-positive T cells were labelled with CellTracker Red (Thermofisher, cat#C34552) at a concentration of 1.5 µM per manufacturer’s instructions. Co-cultures were imaged using an ECHO revolve microscope capturing three fields of view per well. Distance measurements were performed in ImageJ by calibrating images to the embedded scale bar using Analyze > Set Scale. Distances were measured in pixels and converted to micrometers using the calibration factor derived from the scale bar.

#### Sorted subset co-cultures

Conditions for sorting experiments were the same as above, except experiments, T cells were collected by fluorescence automated cell sorted into T cell media using a Sony SH800S with 100 µm chips (Sony, cat#LE-C3210) after overnight digest and CD3 bead-based positive selection. T cells were gated by Lymphocytes/Live/Singlets/CD3+ in bulk sort experiments or Lymphocytes/Live/Singlets/CD4+ or /CD8b+ in sorted subset experiments. In these cases, organoids were co-cultured with approximately 5,000 CD3+ cells, 4,000 CD4+ cells, or 333-500 CD8+ T cells, with the latter two values being based on the relative frequency of T cells in the fetal SI of the initial 5,000 cell inputs. Digested cells were stained using Zombie Viability Dye (1:500, Zombie Violet or Aqua; BioLegend, cat #423113 and 423101), anti-CD3 (1:200; BioLegend, clone SK7 or HIT3a) anti-CD4 (1:200; BioLegend, clone RPA-T4), and anti-CD8b (1:200; BioLegend, clone S21011A).

#### Dual Matrigel^®^ dome organoid cultures

Fetal SI organoids were plated in 24-well plates (CytoOne, cat#CC7682-7524) to allow for two Matrigel^®^ domes per well. Organoids were co-cultured with fetal SI T cells in one dome. In the adjacent dome in the same well, organoids were plated without T cells. For comparison, wells containing no T cells in either dome were included. Cultures were maintained in 500 µl OGM(-Y) per well.

#### Transwell^®^ cultures

Fetal SI organoids were plated in 24-well plates with Transwells*^®^* situated above them (Corning, cat#3413; 0.4 µm pore size). Transwells*^®^* were placed atop 3D-printed O-rings to prevent Transwells*^®^* from pushing down on Matrigel^®^ domes while allowing pore submersion in 500 µl OGM(-Y). Transwells*^®^*were seeded with 5,000 fetal SI T cells in 130 µl T cell media.

### Organoid enumeration

Organoid cultures were imaged via brightfield microscopy using the ECHO Revolve microscope or the Molecular Devices ImageXpress Micro4. For images taken on the ECHO, approximately 20 images through all focal planes were manually captured and stacked into a singular 2D plane via Adobe Photoshop. For images taken on the ImageXpress Micro4, Z-planes were captured and were computationally processed into one 2D image via a minimum projection method. Organoid number and bud length/diameter were manually tallied in a blinded manner. Organoid number is presented as fold change 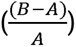 normalized to mock.

### Immunofluorescence

Organoids embedded in Matrigel^®^ were collected on day 6 of culture by loosening the Matrigel^®^ from the plate and collecting it into an Eppendorf tube with 1 mL Advanced DMEM/F12. Matrigel^®^ was dissolved with gentle rocking (∼60-70 rpm) on ice for 1 hour. Whole organoids were pelleted 800xg/5 min/4°C and fixed in 4% paraformaldehyde (Electron Microscopy Sciences, cat #15710) for 30 min on ice. Fixed organoids were then collected by centrifugation at 800 xg/5 min/4°C and resuspended in 1 mL 1X PBS. Organoids were stored at 4°C until staining. Staining was performed via whole-mount methodology as previously described^43^. Organoids were stained using primary antibodies for CHGA (1:100; Agilent DAKO cat#M086901-2) and MUC2 (1:50; Santa Cruz, cat#sc-7314) and species-compatible secondaries (1:400; Invitrogen, cat#A31570 (AF555)) with DAPI (1:2000; Invitrogen D1306) in the final wash.

For MUC2, organoids were subjected to antigen retrieval as follows: fixed organoids were treated with a citrate-based buffer (43.6 mM citric acid, 239.1 mM trisodium citrate dihydrate, 19.8 mM Tris in MilliQ water; pH = 5.7)^44^. Prior to use, the buffer was supplemented with 0.054% Triton X-100. Organoids were resuspended in citrate buffer and boiled for 30 minutes, with buffer replacement from evaporation as necessary. Retrieved samples were then cooled to room temperate before washing once with MilliQ water and proceeding to staining. Stained organoids were mounted on slides using VECTASHIELD Antifade Mounting Medium (Vector Laboraties cat#H-1000-10) and cover slips. Slides imaged via epi-fluorescent microscopy on the ECHO Revolve.

Quantitative analysis of images was performed using a custom automated pipeline implemented in ImageJ/Fiji, based on a three-mask thresholding strategy designed to separately quantify specific signal from background autofluorescence and to exclude non-tissue regions. The fluorescence channel corresponding to the protein of interest was converted to 8-bit grayscale and analyzed using a fixed intensity threshold (20–255). To reduce high-frequency noise and minor intensity fluctuations, a Gaussian blur (σ = 3 pixels) was applied uniformly to all images before mask generation. All intensity measurements were performed on the unblurred original images to preserve true fluorescence values. Bright mucin-associated fluorescence was identified using a high-intensity threshold (pixel values ≥ 20). Tissue autofluorescence was isolated using a narrow low-intensity threshold window (pixel values 3-4). Pixels with two to near zero intensity (pure black areas corresponding to empty space) were excluded from all measurements and were not included in either signal or background masks. For intensity measurements, binary signal and background masks were converted into selections and overlaid onto the corresponding original (unblurred) images. For each image, these metrics were extracted separately for signal and background regions. The corrected Integrated Fluorescence Intensity was calculated using total fluorescence pixel intensity after subtracting the estimated contribution of background autofluorescence. Due to unequal organoid numbers and resulting differences in the number of fields of view (FOVs) collected, analyses were performed using the lowest number of FOVs per donor condition. FOVs were randomly selected in a blinded manner to avoid bias.

### Bulk RNA sequencing and analysis

On day 3 or 6 of culture, organoids were collected into an Eppendorf tube using 1 mL Advanced DMEM/F12. Cell pellets were prepared via centrifugation at 800xg/5 min/4°C and were stored at -80°C until RNA extraction. RNA extraction was performed according to manufacturer’s protocol using the RNeasy Plus Micro Kit (Qiagen, cat#74034), including the DNase I treatment step. RNA concentration and quality were evaluated by the Agilent 2100 Bioanalyzer at the Yale Center for Genome Analysis (YCGA). cDNA libraries for polyA mRNA were prepared by YCGA and quality-control of library size and concentration were evaluated with the Agilent D1000 ScreenTape system. Day 3 libraries were prepared using a low-input preparation method at YCGA, whereas day 6 libraries were prepared as standard input preparations, precluding direct comparison between these two time points in subsequent analysis. Libraries were sequenced on the Illumina NovaSeq with a minimum of 25x10^6^ reads per sample.

Raw sequencing reads were assessed for quality using FastQC (v0.12.1)^45^. Adapter sequences and low-quality bases were removed using Trimmomatic (v0.39)^46^. Trimmed reads were aligned to the human reference genome (GRCh38/hg38) via HISAT2 (v2.2.1)^47^, and post-alignment quality metrics were assessed with Picard (v2.25.6). Gene-level quantification was performed using StringTie (v2.2.1)^48^ in reference-guided mode with Ensembl GRCh38 gene annotations. Individual sample counts were compiled into a unified gene count matrix for differential expression analysis. Gene count matrices were loaded into R (v2024.12.1), and size factors were determined for each to normalize for variation in sequencing depth, library size, etc. Genes with less than 10 counts across all samples were removed. Differential gene expression analysis was conducted using DESeq2 ^49^ after adaptive shrinkage estimator (ashr) to reduce noise from low-count genes^50^. Genes were considered differentially expressed at an adjusted p-value threshold of 0.05 (Benjamini-Hochberg correction). Analysis was restricted to protein-coding genes as annotated in org.Hs.eg.db. Pathway analysis was conducted using Gene Set Enrichment Analysis (GSEA)^51^ with Gene Ontology for biological processes with a significance threshold of 0.05. Data visualization and plotting were performed using ggplot2, pHeatmap, Enhanced Volcano, ggrepel, dplyr, tidyr, and plotly R packages^52–58^. Computational deconvolution was performed via Bisque^22^.

Independent sequencing batches were integrated using a multi-step batch correction strategy that leveraged mock-cultured organoid samples as biological anchors. Raw gene counts were first filtered using a condition-aware approach: genes were retained if they showed sufficient expression (counts >10 in at least 2 samples) in any experimental condition, preserving genes specifically induced by fetal or adult T cells that might be absent in other conditions. For batch correction, we employed ComBat-seq on the filtered raw count matrix, using the sva package^59^. Mock organoid conditions, which were present in both sequencing batches, served as biological reference points to estimate technical variation between batches. The ComBat-seq algorithm was applied with full modeling (full_mod = TRUE) to preserve the biological signal from T cell age groups while removing batch-associated technical variation. To validate the effectiveness of batch correction, we performed principal component analysis on rlog-transformed counts before and after correction using DESeq2, confirming that batch-associated clustering was eliminated while biological groupings remained distinct.

### Single-cell RNA sequencing and analysis

In **Figure 3**, organoids were grown with or without fetal SI T cells in OGM(-Y) for 3 days. In **Figure 4**, cultures were grown for 2 days with OGM(-Y), then the media was substituted for IntestiCult Human Organoid Differentiation Media (StemCell Technologies, cat #100-0214) containing 5 µM DAPT (StemCell Technologies, cat #72082) in a ratio of 3:1 supplement:basal for 2 days. On day 4, media was substituted fully to a 1:1 ratio of supplement:basal for the final 2 days. Organoids were collected on day 3 (growth phase scRNA-seq) or day 6 (differentiation phase scRNA-seq) in Cell Recovery Solution (Corning, cat #354253) into a conical tube. Organoids gently rocking (∼60-70 rpm) at 4°C for 15 mins then pelleted by centrifugation at 500xg/5 min/4°C. Organoids were resuspended in TrypLE Express (Gibco, cat#12605-010) and incubated at 37°C/5% CO_2_ with manual trituration every 5 mins for 20 mins total. Dissociated cells were centrifuged at 300xg/10 min/4°C and were resuspended in 0.1% BSA in 1X PBS. Viability was assessed using Trypan blue staining, and samples with viability <80% underwent dead cell removal using MS columns. Cells were resuspended in RPMI and counted, ensuring viability >80%. If viability was under this threshold, another dead cell removal was performed.

Single-cell transcript count matrices generated by Cell Ranger (10X Genomics) were used for the OGM and ODM organoid datasets. All downstream preprocessing and clustering analyses were performed using the Scanpy (v1.9.2) package^60^. For each dataset, cells were filtered to retain those with more than 500 detected genes and less than 25% mitochondrial gene expression, and genes expressed in fewer than three cells were excluded. Doublets were identified using Scrublet (v0.2.3), and doublet scores were calculated for each cell^61^. Gene expression values were normalized on a per-cell basis and log-transformed. Highly variable genes were subsequently identified using scanpy.pp.highly_variable_genes with default parameters. To mitigate technical effects, the percentage of mitochondrial gene expression and unique molecular identifier (UMI) counts were regressed out using scanpy.pp.regress_out, followed by data scaling.

Batch correction across samples was performed using BBKNN (v1.5.1) on the first 50 principal components^62^. Dimensionality reduction and Leiden clustering were carried out using the resulting highly variable genes, and cells were visualized using Uniform Manifold Approximation and Projection (UMAP). Clusters exhibiting the highest predicted doublet scores were excluded from downstream analyses. Cell lineages were manually annotated based on established marker genes reported in the literature, together with differentially expressed genes identified for each cluster using scanpy.tl.rank_genes_groups with the Wilcoxon rank-sum test. Relative T cell population frequencies were evaluated by single-cell annotation using variational inference (scANVI)^63^.

### Spatial transcriptomics and analysis

#### Sample preparation

Formalin fixed paraffin embedded tissue blocks were sectioned as tissue microarray (TMA) cores using a tissue arrayer (Beecher Instruments) onto Xenium slides included in the Xenium Prime 5K Human Pan Tissue & Pathways Assay Kit (10X Genomics, cat#1000671) at the Yale Histology Core. Tissue microarrays were processed by Xenium Prime 5K at the Yale Center for Genome Analysis within 48 hours of placement.

#### Processing, batch correction, and analysis

The Xenium spatial transcriptomics data were processed using the Xenium Ranger (v4.0.0) pipeline for read alignment and cell segmentation, employing the default nuclei-based segmentation workflow. Tissues from the TMA were identified by manually delineating tissue boundaries using selection tools in Xenium Explorer (v3.2). Tissue identifiers and corresponding x–y coordinates were exported as .csv files for each slide.

Cell-by-gene count matrices generated by Xenium Explorer were converted into Scanpy-compatible .h5ad objects. Individual tissues were then separated into distinct .h5ad files based on their corresponding tissue IDs and spatial coordinates. Cells with fewer than 10 detected genes or fewer than 20 transcripts were filtered out, and genes expressed in fewer than 10 cells were removed. Gene expression values were normalized on a per-cell basis and log-transformed. Highly variable genes were identified using scanpy.pp.highly_variable_genes with the cell_ranger flavor, selecting the top 2,000 genes (n_top_genes = 2000) and default parameters. To account for technical variation, UMI counts were regressed out using scanpy.pp.regress_out, followed by data scaling.

Batch correction across samples was performed using Harmony (harmonypy) (v.0.0.10) on the first 50 principal components^64^. Dimensionality reduction and Leiden clustering were conducted using the highly variable genes, and cells were visualized using UMAP. Clusters with the highest predicted doublet scores were excluded from downstream analyses. Cell lineages were manually annotated based on established marker genes reported in the literature, in combination with differentially expressed genes identified for each cluster using scanpy.tl.rank_genes_groups with the Wilcoxon rank-sum test.To ensure the integrity of the cell segmentation and mitigate artifacts such as segmentation leakage or cellular fusion, we calculated the mutually exclusive correlation ratio (MECR)^65^. This check involved quantifying the co-expression of canonical markers known to be mutually exclusive across distinct cell lineages within individual segmented cell boundaries. The MECR score was calculated using a combination of universal cell-type markers and specialized metrics designed to address segmentation challenges unique to the intestinal epithelium. We prioritized the identification of intraepithelial lymphocytes, which frequently reside in such close physical proximity to epithelial cells that they are prone to mis-segmentation. The MECR gene set was adapted from other spatial transcriptomics studies^66^. Samples displaying high MECR were flagged for manual review to ensure that observed spatial interactions were not biased by segmentation inaccuracies.

Following successful QC, spatial interaction metrics were quantified using Squidpy (v1.6.5). Euclidean distances between T-cell coordinates and the nearest neighbor within defined epithelial subsets (Stem, Crypt, and All Epithelial) were computed using a pairwise distance matrix. A spatial proximity threshold of 50 µm was applied to determine the percentage of T-cells residing within the immediate epithelial microenvironment.

#### Global spatial environment mapping

To objectively identify spatially conserved tissue microenvironments (niches) in an unbiased manner, we employed an unsupervised spatial clustering approach. To mitigate batch effects and tissue-specific variance, each tissue section was processed independently. For each slide, we constructed a spatial neighbor graph using the **squidpy** framework, defining spatial connectivities via a k-nearest neighbor (kNN) approach (k=6) based on tissue-specific spatial coordinates. This connectivity threshold was selected to capture the immediate paracrine neighborhood of each cell while maintaining sensitivity to local tissue topology. Following graph construction, we applied the **Leiden algorithm** to the spatial connectivity matrix to identify locally cohesive clusters. This segmentation approach allowed for the unbiased partitioning of the tissue into discrete, spatially-informed niches. Each identified niche was assigned a unique identifier by concatenating the sample ID and the Leiden cluster label.

To validate the biological relevance of the identified spatial niches, we performed manual anatomical verification on a representative subset of tissues (n=3 samples per condition). We systematically overlaid niche identifiers onto high-resolution tissue images to assess their correspondence with known anatomical landmarks, such as the crypt-villus axis and the lamina propria. Clusters that did not demonstrate clear spatial cohesion or anatomical interpretability were excluded from downstream analyses.

#### Cell-cell communication inference

To infer potential ligand-receptor interactions with spatial specificity, we utilized a microenvironment-aware implementation of CellPhoneDB v5.^67^ We defined spatial niches using Leiden clustering on the spatial connectivity graph (as described above) and mapped each cell to its corresponding microenvironment. This metadata was provided to CellPhoneDB via a microenvironment-specification file, which restricted the permutation test (1,000 permutations) to interactions occurring within the same spatial niche to ensure that reported signaling events were statistically significant within their local spatial context, effectively excluding signaling pairs that were not co-localized within the same tissue microenvironment. Statistical significance was defined as p < 0.05 with a minimum gene expression threshold of 10% across the cell populations. From the resulting interactome, we focused our downstream analysis on the bi-directional crosstalk between T cell subtypes and ISCs.

### Luminex

Supernatant from cultures were pooled between technical replicate wells and frozen at -80°C. Samples were sent to Eve Technologies and were analyzed using the Human Cytokine/Chemokine 96-Plex Discovery Assay® (HD96) for data in **Figure 6** and **Figure S5** or the Human Metabolic Hormone 12-Plex Discovery Assay® (HDMET12) in **Figure 4**. For HD96, quantile normalization and log2 transformation were performed, followed by principal component analysis. Cytokines/chemokines that were undetectable in all samples were removed from analysis, and those that were undetectable in only some conditions were imputed as half of the lowest detected concentration in experimental samples. Comparisons between fetal and adult co-cultures were assessed after using removeBatchEffect in the limma package to remove technical variation from sequencing batch effects^68^. Pairwise differential expression was performed between conditions, where p<0.05 and |L2FC|>1.5 or >2 were considered significant as noted.

### Conditioned media cultures

#### Thawed and unconcentrated conditioned media

Media from day 3 of fetal SI T cell-organoid co-cultures was collected from triplicate wells and stored at -80°C. After thawing the conditioned media from mock or +T cell cultures on ice, it was diluted by approximately half (∼0.8:1 conditioned media:OGM(-Y)) and then used as culture media for newly passaged fetal organoids. Conditioned media cultures were ended on day 3.

#### Thawed and concentrated conditioned media

On day 2 of cultures, media from fetal SI T cell-organoid co-cultures was pooled from triplicate wells and stored at -80°C. After thawing the media on ice, the media was concentrated using 3k MWCO Omega concentrator columns (Pall, cat#MAP003C37) to a final volume of 50 µl according to the manufacturer’s protocol. Media was reconstituted at a ratio of 1:15 concentrate:fresh using OGM(-Y). Organoids were grown in concentrated conditioned media for 3 days.

#### Freshly concentrated conditioned media (without freeze-thaw)

Fetal organoids were co-cultured with or without fetal SI T cells after CD3 positive selection described above in OGM(-Y). Media (700 µl, pooled from triplicate wells) was collected from the mock and T cell co-cultures on day 2, and media was concentrated using 3k MWCO Omega concentrator columns to a final volume of 50 µl. At this time, the mock or T cell co-cultures were given fresh OGM(-Y) and returned to 37°C/5% CO_2_ to collect again 2 days later. The concentrated media was reconstituted at volume with fresh OGM(-Y) at a ratio of 1:15 concentrate:fresh. Fetal organoids were passaged as described above and plated in triplicate. Organoids were cultured with 250 µl/well of the reconstituted conditioned media. This process was repeated once more 2 days later to replenish reconstituted conditioned media. Cultures were ended on day 4.

### Statistical analysis and data availability

Data were analyzed using Prism 10 and assessed for normality by the Shapiro-Wilk test. Parametric data were compared by unpaired t-test with Welch’s correction or one-way analysis of variance (ANOVA) with Tukey’s post-hoc. Nonparametric data were analyzed via Mann-Whitney or Kruskal-Wallis with Dunn’s multiple comparisons test. Differences were considered statistically significant with p<0.05 with specific tests used denoted in each figure legend. In spatial transcriptomic distance comparisons, all p values were adjusted using the Bonferroni correction (p_adj = p x n, where n is the number of experimental conditions) to mitigate the risk of type I error inflation. Differences were considered statistically significant at an adjusted threshold of p < 0.05.All data are reported in this paper or are available at NCBI GEO under accession XX (to be made available upon publication). Code is available at Github XX (to be made available upon publication).

## CONTRIBUTION STATEMENT

Conceptualization: LK, MSS Methodology: LK, MSS

Investigation: MSS, KG, LP, WG, HY, WW, DL-L, KS, EGGS, JL

Visualization: MSS, LP Funding acquisition: LK, MSS Supervision: LK

Writing – original draft: MSS

Writing – reviewing/editing: MSS, LP, KG, WG, HY, WW, DL-L, KS, EGGS, JL, DZ, DTB, GT, LK

## ACKNOWLEDGMENTS

We thank the Yale Center for Research Computing, the Yale Flow Cytometry core (supported in part by an NCI Cancer Center Support Grant # NIH P30 CA016359), the Yale Center for Genome Analysis core (supported by the National Institute of General Medical Sciences of the NIH under Award #1S10OD030363-01A1), and the Yale Pathology Tissue Services Shared Resource for histology services (supported in part by the National Cancer Institute of the National Institutes of Health under Award #P30CA016359). We would also like to thank Dr. Upasana Das Adhikari and Dr. Oluwabunmi Olaloye for helpful discussions.

## REFERENCES

1. Sato, T., Vries, R.G., Snippert, H.J., van de Wetering, M., Barker, N., Stange, D.E., van Es, J.H., Abo, A., Kujala, P., Peters, P.J., et al. (2009). Single Lgr5 stem cells build crypt-villus structures in vitro without a mesenchymal niche. Nature 459. 10.1038/nature07935.

2. Gehart, H., Clevers, H., Gehart, H., and Clevers, H. (2018). Tales from the crypt: new insights into intestinal stem cells. Nature Reviews Gastroenterology & Hepatology 16. 10.1038/s41575-018-0081-y.

3. Clevers, H., Loh, K.M., and Nusse, R. (2014). An integral program for tissue renewal and regeneration: Wnt signaling and stem cell control. Science 346. 10.1126/science.1248012.

4. Demitrack, E.S., and Samuelson, L.C. (2016). Notch regulation of gastrointestinal stem cells. The Journal of Physiology 594. 10.1113/JP271667.

5. Holloway, E.M., Czerwinski, M., Tsai, Y.-H., Wu, J.H., Wu, A., Childs, C.J., Walton, K.D., Sweet, C.W., Yu, Q., Glass, I., et al. (2021). Mapping Development of the Human Intestinal Niche at Single-Cell Resolution. Cell Stem Cell 28. 10.1016/j.stem.2020.11.008.

6. Gu, W., Eke, C., Santiago, E., Olaloye, O., and Konnikova, L. (2024). Single-cell atlas of the small intestine throughout the human lifespan demonstrates unique features of fetal immune cells. Mucosal Immunology 17, 599–617. 10.1016/j.mucimm.2024.03.011.

7. Elmentaite, R., Kumasaka, N., Roberts, K., Fleming, A., Dann, E., King, H., Kleshchevnikov, V., Dabrowska, M., Pritchard, S., Bolt, L., et al. (2021). Cells of the human intestinal tract mapped across space and time. Nature 597. 10.1038/s41586-021-03852-1.

8. Park, J.-E., Jardine, L., Gottgens, B., Teichmann, S.A., and Haniffa, M. (2020). Prenatal development of human immunity. Science 368. 10.1126/science.aaz9330.

9. Barnes, J.L., Yoshida, M., He, P., Worlock, K.B., Lindeboom, R.G.H., Suo, C., Pett, J.P., Wilbrey-Clark, A., Dann, E., Mamanova, L., et al. (2023). Early human lung immune cell development and its role in epithelial cell fate. Science Immunology 8. 10.1126/sciimmunol.adf9988.

10. Stras, S., Werner, L., Toothaker, J., Olaloye, O., Oldham, A., McCourt, C., Lee, Y., Rechavi, E., Shouval, D., and Konnikova, L. (2019). Maturation of the Human Intestinal Immune System Occurs Early in Fetal Development. Developmental Cell 51, 357–373.e355. 10.1016/j.devcel.2019.09.008.

11. Feyaerts, D., Urbschat, C., Gaudillière, B., and Stelzer, I. (2022). Establishment of tissue-resident immune populations in the fetus. Seminars in immunopathology 44, 747–766. 10.1007/s00281-022-00931-x.

12. Schreurs, R., Baumdick, M., Sagebiel, A., Kaufmann, M., Mokry, M., Klarenbeek, P., Schaltenberg, N., Steinert, F., van Rijn, J., Drewniak, A., et al. (2019). Human Fetal TNF-α-Cytokine-Producing CD4+ Effector Memory T Cells Promote Intestinal Development and Mediate Inflammation Early in Life. Immunity 50, 462–476.e468. 10.1016/j.immuni.2018.12.010.

13. Egozi, A., Olaloye, O., Werner, L., Silva, T., McCourt, B., Pierce, R., An, X., Wang, F., Chen, K., Pober, J., et al. (2023). Single-cell atlas of the human neonatal small intestine affected by necrotizing enterocolitis. PLoS Biology 21. 10.1371/journal.pbio.3002124.

14. He, S., Luo, C.-L., Luo, T., Chen, H.-T., Zhang, S.-F., Jiang, J.-X., Wang, X.-Y., Ma, D., Zhao, S.-L., Xu, A.-Y., et al. (2025). Systemic immune activity occurs during human immune system maturation. Cell 188. 10.1016/j.cell.2025.10.003.

15. Mahapatro, M., Erkert, L., and Becker, C. (2021). Cytokine-Mediated Crosstalk between Immune Cells and Epithelial Cells in the Gut. Cells 10. 10.3390/cells10010111.

16. Haber, A., Biton, M., Rogel, N., Herbst, R., Shekhar, K., Smillie, C., Burgin, G., Delorey, T., Howitt, M., Katz, Y., et al. (2017). A single-cell survey of the small intestinal epithelium. Nature 551, 333–339. doi:10.1038/nature24489.

17. Recaldin, T., Steinacher, L., Gjeta, B., Harter, M.F., Adam, L., Kromer, K., Mendes, M.P., Bellavista, M., Nikolaev, M., Lazzaroni, G., et al. (2024). Human organoids with an autologous tissue-resident immune compartment. Nature. 10.1038/s41586-024-07791-5.

18. Noel, G., Baetz, N.W., Staab, J.F., Donowitz, M., Kovbasnjuk, O., Pasetti, M.F., and Zachos, N.C. (2017). A primary human macrophage-enteroid co-culture model to investigate mucosal gut physiology and host-pathogen interactions. Scientific Reports 7, 45270. 10.1038/srep45270.

19. Sharafian, Z., Littlejohn, P.T., Michalski, C., Sousa, J.A., Cheung, J., Hill, M., Piper, H., Jacobson, K., Lavoie, P.M., Allaire, J.M., and Vallance, B.A. (2025). Crosstalk with infant-derived Th17 cells, as well as exposure to IL-22 promotes maturation of intestinal epithelial cells in an enteroid model. Frontiers in Immunology 16. 10.3389/fimmu.2025.1582688.

20. Santos, A.J.M., van Unen, V., Lin, Z., Chirieleison, S.M., Ha, N., Batish, A., Chan, J.E., Cedano, J., Zhang, E.T., Mu, Q., et al. (2024). A human autoimmune organoid model reveals IL-7 function in coeliac disease. Nature 2024 632:8024 632. 10.1038/s41586-024-07716-2.

21. Hansen, S.L., Larsen, H.L., Pikkupeura, L.M., Maciag, G., Guiu, J., Iris Müller, Clement, D.L., Mueller, C., Johansen, J.V., Helin, K., et al. (2023). An organoid-based CRISPR-Cas9 screen for regulators of intestinal epithelial maturation and cell fate. Science Advances 9. 10.1126/sciadv.adg4055.

22. Jew, B., Alvarez, M., Rahmani, E., Miao, Z., Ko, A., Garske, K.M., Sul, J.H., Pietiläinen, K.H., Pajukanta, P., Halperin, E., et al. (2020). Accurate estimation of cell composition in bulk expression through robust integration of single-cell information. Nature Communications 11. 10.1038/s41467-020-15816-6.

23. Yang, L., Wang, X., Zhou, X., Chen, H., Song, S., Deng, L., Yao, Y., Yin, X., Yang, L., Wang, X., et al. (2025). A tunable human intestinal organoid system achieves controlled balance between self-renewal and differentiation. Nature Communications 16. 10.1038/s41467-024-55567-2.

24. Egozi, A., Llivichuzhca-Loja, D., McCourt, B.T., Bahar Halpern, K., Farack, L., An, X., Wang, F., Chen, K., Konnikova, L., Itzkovitz, S., et al. (2021). Insulin is expressed by enteroendocrine cells during human fetal development. Nature Medicine 27. 10.1038/s41591-021-01586-1.

25. Gribble, F., and Reimann, F. (2016). Enteroendocrine Cells: Chemosensors in the Intestinal Epithelium. Annual Review of Physiology 78. 10.1146/annurev-physiol-021115-105439.

26. Biton, M., Haber, A.L., Rogel, N., Burgin, G., Beyaz, S., Schnell, A., Ashenberg, O., Su, C.W., Smillie, C., Shekhar, K., et al. (2018). T helper cell cytokines modulate intestinal stem cell renewal and differentiation. Cell 175. 10.1016/j.cell.2018.10.008.

27. Fawkner-Corbett, D., Antanaviciute, A., Parikh, K., Jagielowicz, M., Gerós, A., Gupta, T., Ashley, N., Khamis, D., Fowler, D., Morrissey, E., et al. (2021). Spatiotemporal analysis of human intestinal development at single-cell resolution. Cell 184, 810–826.e823. 10.1016/j.cell.2020.12.016.

28. Olaloye, O., Liu, P., Toothaker, J., McCourt, B., McCourt, C., Xiao, J., Prochaska, E., Shaffer, S., Werner, L., Gringauz, J., et al. (2021). CD16+CD163+ monocytes traffic to sites of inflammation during necrotizing enterocolitis in premature infants. The Journal of Experimental Medicine 218. 10.1084/jem.20200344.

29. Denning, T., Bhatia, A., Kane, A., Patel, R., and Denning, P. (2017). Pathogenesis of NEC: Role of the innate and adaptive immune response. Seminars in Perinatology 41. 10.1053/j.semperi.2016.09.014.

30. Demers-Mathieu, V. (2022). The immature intestinal epithelial cells in preterm infants play a role in the necrotizing enterocolitis pathogenesis: A review. Health Sciences Review 4. 10.1016/j.hsr.2022.100033.

31. Zhou, Q., Niño, D.F., Yamaguchi, Y., Wang, S., Fulton, W.B., Jia, H., Lu, P., Prindle Jr., T., Pamies, D., Morris, M., et al. (2021). Necrotizing enterocolitis induces T lymphocyte–mediated injury in the developing mammalian brain. Science Translational Medicine 13. 10.1126/scitranslmed.aay6621.

32. Zhang, Z., Lee, D., Liu, L., Xiong, Y., Lee, C., Kim, J.-E., Chusilp, S., Lau, E., Tian, Y., Feizi, M., et al. (2025). Impairment of stromal-epithelial regenerative cross-talk in Hirschsprung disease primes for the progression to enterocolitis. Science Translational Medicine 17. 10.1126/scitranslmed.adp4679.

33. Weaver, L., Austin, S., and Cole, T. (1991). Small intestinal length: a factor essential for gut adaptation. Gut 32. 10.1136/gut.32.11.1321.

34. Marnerides, A., Ghazi, S., Sundberg, A., Papadogiannakis, N., and Andreas Marnerides, S.G., Anders Sundberg, Nikos Papadogiannakis (2012). Development of Fetal Intestinal Length during 2nd-Trimester in Normal and Pathologic Pregnancies. Pediatric and Developmental Pathology 15. 10.2350/11-07-1057-OA.1.

35. Connors, T.J., Matsumoto, R., Verma, S., Szabo, P.A., Guyer, R., Gray, J., Wang, Z., Thapa, P., Dogra, P., Poon, M.M.L., et al. (2023). Site-specific development and progressive maturation of human tissue-resident memory T cells over infancy and childhood. Immunity 56. 10.1016/j.immuni.2023.06.008.

36. Halkias, J., Rackaityte, E., Hillman, S.L., Aran, D., Mendoza, V.F., Marshall, L.R., MacKenzie, T.C., and Burt, T.D. (2019). CD161 contributes to prenatal immune suppression of IFN-γ–producing PLZF+ T cells. The Journal of Clinical Investigation 129. 10.1172/JCI125957.

37. Mitra, A., Shanthalingam, S., Sherman, H.L., Singh, K., Canakci, M., Torres, J.A., Lawlor, R., Ran, Y., Golde, T.E., Miele, L., et al. (2020). CD28 Signaling Drives Notch Ligand Expression on CD4 T Cells. Frontiers in Immunology 11. 10.3389/fimmu.2020.00735.

38. Teixé, T., Nieto-Blanco, P., Vilella, R., Engel, P., Reina, M., and Espel, E. (2008). Syndecan-2 and -4 expressed on activated primary human CD4+ lymphocytes can regulate T cell activation. Molecular Immunology 45. 10.1016/j.molimm.2008.01.033.

39. Li, N., van Unen, V., Abdelaal, T., Guo, N., Kasatskaya, S., Ladell, K., McLaren, J., Egorov, E., Izraelson, M., Chuva de Sousa Lopes, S., et al. (2019). Memory CD4+ T cells are generated in the human fetal intestine. Nature immunology 20. 10.1038/s41590-018-0294-9.

40. Wosen, J.E., Ilstad-Minnihan, A., Co, J.Y., Jiang, W., Mukhopadhyay, D., Fernandez-Becker, N.Q., Kuo, C.J., Amieva, M.R., and Mellins, E.D. (2019). Human Intestinal Enteroids Model MHC-II in the Gut Epithelium. Frontiers in Immunology 10. 10.3389/fimmu.2019.01970.

41. Oh, C. (2023). Jejunoileal Atresia: A Contemporary Review. Advances in Pediatric Surgery 29. 10.13029/aps.2023.29.2.89.

42. Berseth, C.L. (1994). Gut Motility and the Pathogenesis of Necrotizing Enterocolitis. Clinics in Perinatology 21. 10.1016/S0095-5108(18)30345-2.

43. Smith, L., Santiago, E.G., Eke, C., Gu, W., Wang, W., Llivichuzhca-Loja, D., Kehoe, T., St Denis, K., Strine, M., Taylor, S., et al. (2024). Human Milk Supports Robust Intestinal Organoid Growth, Differentiation, and Homeostatic Cytokine Production. Gastro Hep Advances 3, 1030–1042. 10.1016/j.gastha.2024.07.007.

44. Toulehohoun, A.G., Bouzin, C., Daumerie, A., Maccioni, L., and Stärkel, P. (2025). 3D fluorescence staining and confocal imaging of low amount of intestinal organoids (enteroids): Protocol accessible to all. PLOS ONE 20. 10.1371/journal.pone.0315922.

45. Andrews, S. (2010). FastQC: A Quality Control Tool for High Throughput Sequence Data.

46. Bolger, A., Lohse, M., and Usadel, B. (2014). Trimmomatic: a flexible trimmer for Illumina sequence data - PubMed. Bioinformatics 30. 10.1093/bioinformatics/btu170.

47. Kim, D., Paggi, J., Park, C., Bennett, C., and Salzberg, S. (2019). Graph-based genome alignment and genotyping with HISAT2 and HISAT-genotype. Nature Biotechnology 37. 10.1038/s41587-019-0201-4.

48. Pertea, M., Pertea, G., Antonescu, C., Chang, T., Mendell, J., and Salzberg, S. (2015). StringTie enables improved reconstruction of a transcriptome from RNA-seq reads - PubMed. Nature Biotechnology 33. 10.1038/nbt.3122.

49. Love, M., Huber, W., and Anders, S. (2014). Moderated estimation of fold change and dispersion for RNA-seq data with DESeq2. Genome biology 15. 10.1186/s13059-014-0550-8.

50. Stephens, M. (2017). False discovery rates: a new deal. Biostatistics 18. 10.1093/biostatistics/kxw041.

51. Subramanian, A., Tamayo, P., Mootha, V., Mukherjee, S., Ebert, B., Gillette, M., Paulovich, A., Pomeroy, S., Golub, T., Lander, E., and Mesirov, J. (2005). Gene set enrichment analysis: a knowledge-based approach for interpreting genome-wide expression profiles - PubMed. Proceedings of the National Academy of Sciences 102. 10.1073/pnas.0506580102.

52. Blighe, K., Rana, S., and Lewis, M. (2023). EnhancedVolcano: Publication-ready volcano plots with enhanced colouring and labeling. R package version 1.20.0.

53. Kolde, R. (2019). pheatmap: Pretty Heatmaps. R package version 1.0.12.

54. Slowikowski, K. (2023). ggrepel: Automatically Position Non-Overlapping Text Labels with ’ggplot2’. R package version 0.9.3.

55. Wickham, H., François, R., Henry, L., Müller, K., and Vaughan, D. (2023). dplyr: A Grammar of Data Manipulation. R package version 1.1.2.

56. Wickham, H., Vaughan, D., and Girlich, M. (2023). tidyr: Tidy Messy Data. R package version 1.3.0.

57. Sievert, C. (2020). Interactive Web-Based Data Visualization with R, plotly, and shiny. Chapman and Hall/CRC.

58. Wickham, H. (2016). ggplot2: Elegant Graphics for Data Analysis.

59. Leek, J.T., Johnson, W.E., Parker, H.S., Jaffe, A.E., and Storey, J.D. (2012). The sva package for removing batch effects and other unwanted variation in high-throughput experiments. Bioinformatics 28. 10.1093/bioinformatics/bts034.

60. Wolf, F., Angerer, P., and Theis, F. (2018). SCANPY: large-scale single-cell gene expression data analysis. Genome Biology 19. 10.1186/s13059-017-1382-0.

61. Wolock, S., Lopez, R., and Klein, A. (2019). Scrublet: Computational Identification of Cell Doublets in Single-Cell Transcriptomic Data. Cell Systems 8. 10.1016/j.cels.2018.11.005.

62. Polański, K., Young, M., Miao, Z., Meyer, K., Teichmann, S., and Park, J. (2020). BBKNN: fast batch alignment of single cell transcriptomes. Bioinformatics 36. 10.1093/bioinformatics/btz625.

63. Xu, C., Lopez, R., Mehlman, E., Regier, J., Jordan, M., and Yosef, N. (2021). Probabilistic harmonization and annotation of single-cell transcriptomics data with deep generative models. Molecular Systems Biology 17. 10.15252/msb.20209620.

64. Korsunsky, I., Millard, N., Fan, J., Slowikowski, K., Zhang, F., Wei, K., Baglaenko, Y., Brenner, M., Loh, P., and Raychaudhuri, S. (2019). Fast, sensitive and accurate integration of single-cell data with Harmony. Nature Methods 16. 10.1038/s41592-019-0619-0.

65. Plummer, J.T., Dezem, F.S., Cook, D.P., Park, J., Zhang, L., Liu, Y., Marção, M., DuBose, H., Wani, A., Wise, K., et al. (2025). Standardized metrics for assessment and reproducibility of imaging-based spatial transcriptomics datasets. Nature Biotechnology. 10.1038/s41587-025-02811-9.

66. McGregor, C., Qin, X., Jagielowicz, M., Gupta, T., Yin, Z., Lentsch, V., Fawkner-Corbett, D., Wien Lai, V., Gomez Castro, P., Bridges, E., et al. (2025). Spatial fibroblast niches define Crohn’s fistulae. Nature 649. 10.1038/s41586-025-09744-y.

67. Troulé, K., Petryszak, R., Cakir, B., Cranley, J., Harasty, A., Prete, M., Tuong, Z.K., Teichmann, S.A., Garcia-Alonso, L., Vento-Tormo, R., et al. (2025). CellPhoneDB v5: inferring cell–cell communication from single-cell multiomics data. Nature Protocols 20. 10.1038/s41596-024-01137-1.

68. Ritchie, M., Phipson, B., Wu, D., Hu, Y., Law, C., Shi, W., and Smyth, G. (2015). limma powers differential expression analyses for RNA-sequencing and microarray studies. Nucleic Acids Research 43. 10.1093/nar/gkv007.

