## Supplementary material for "Early-life mucosal T cells direct intestinal stem cell fate via a coordinated developmental program": supply fig

**Supplemental Figure 1.**

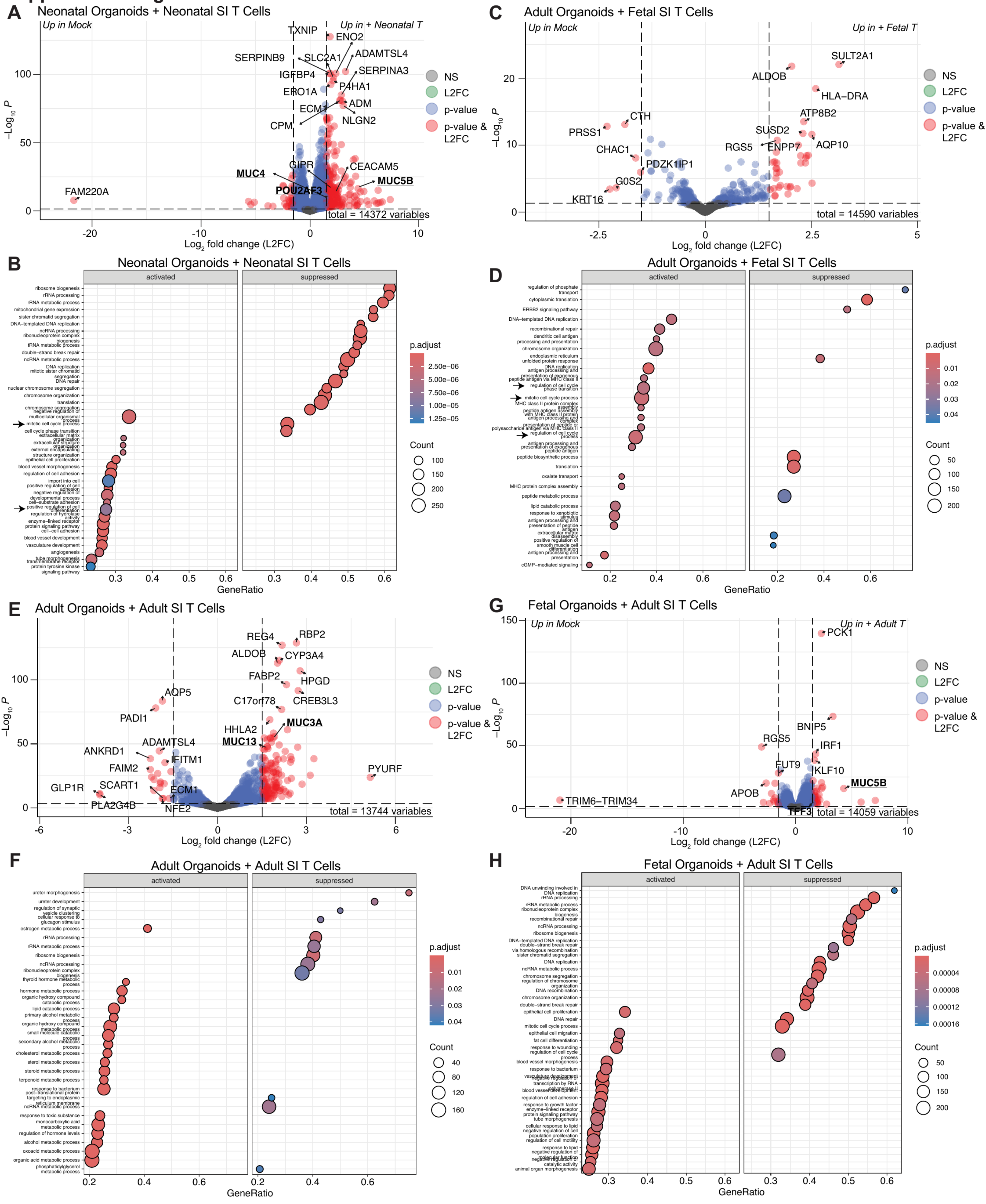

#### Supplemental Figure 2.

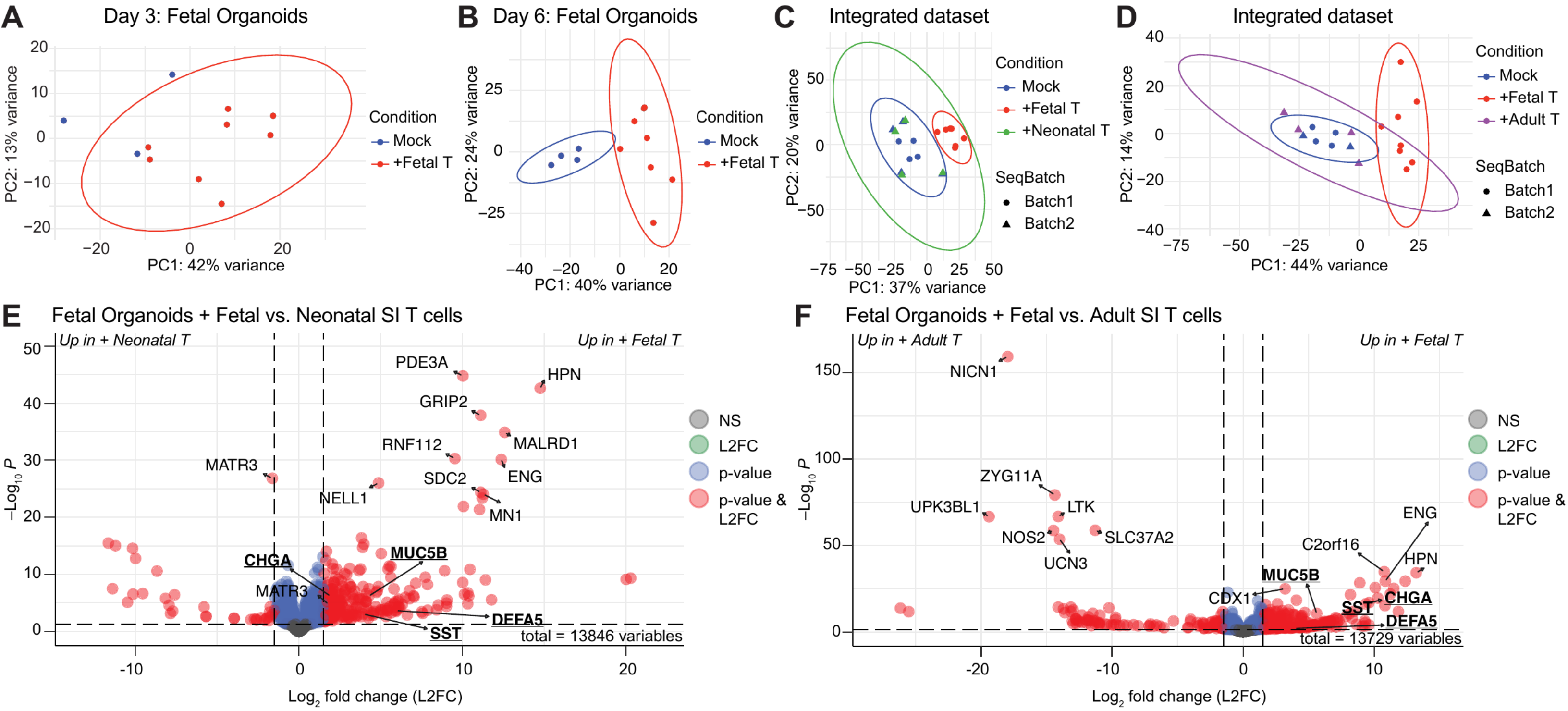

Supplemental Figure 3.

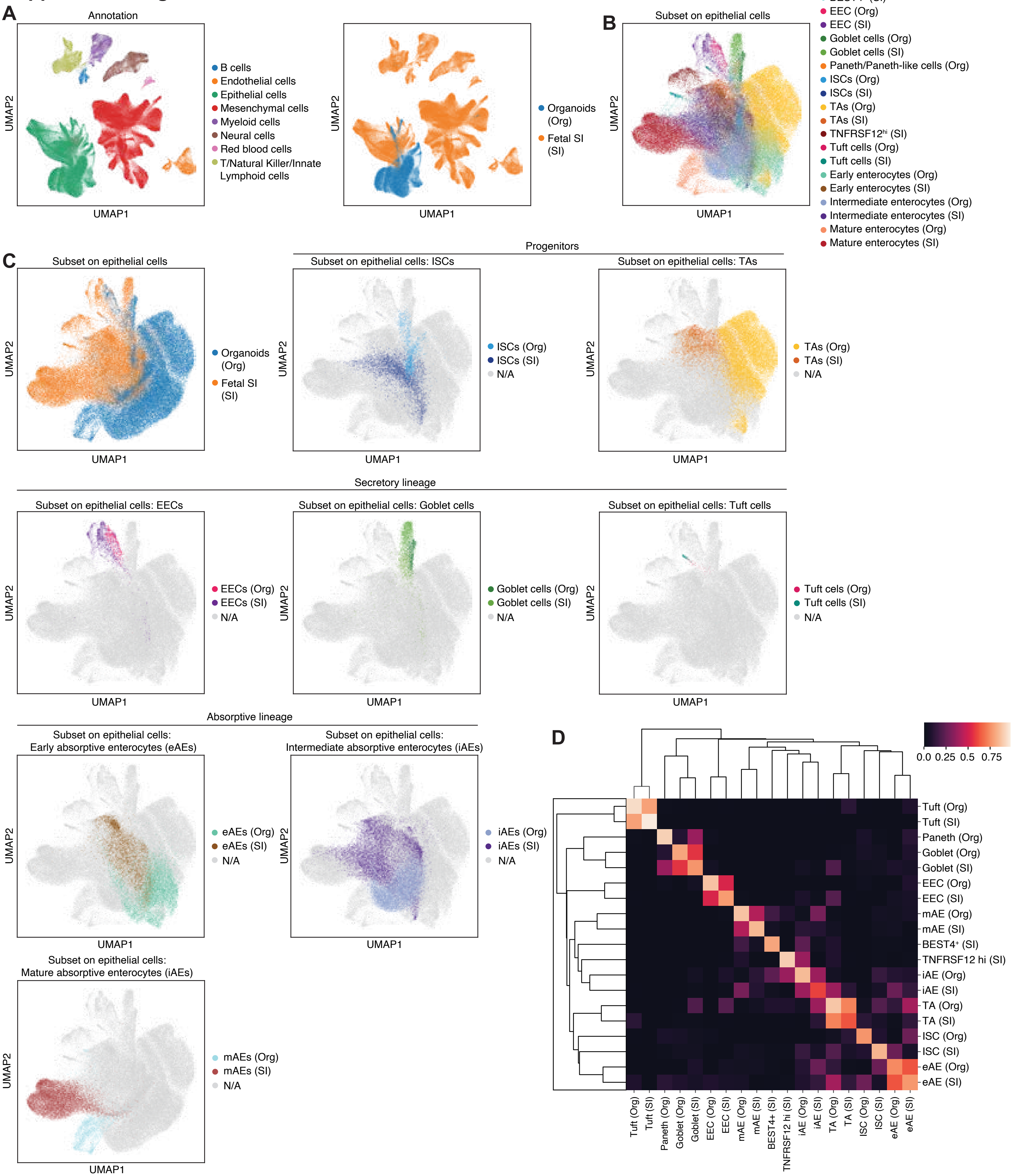

### Supplemental Figure 4.

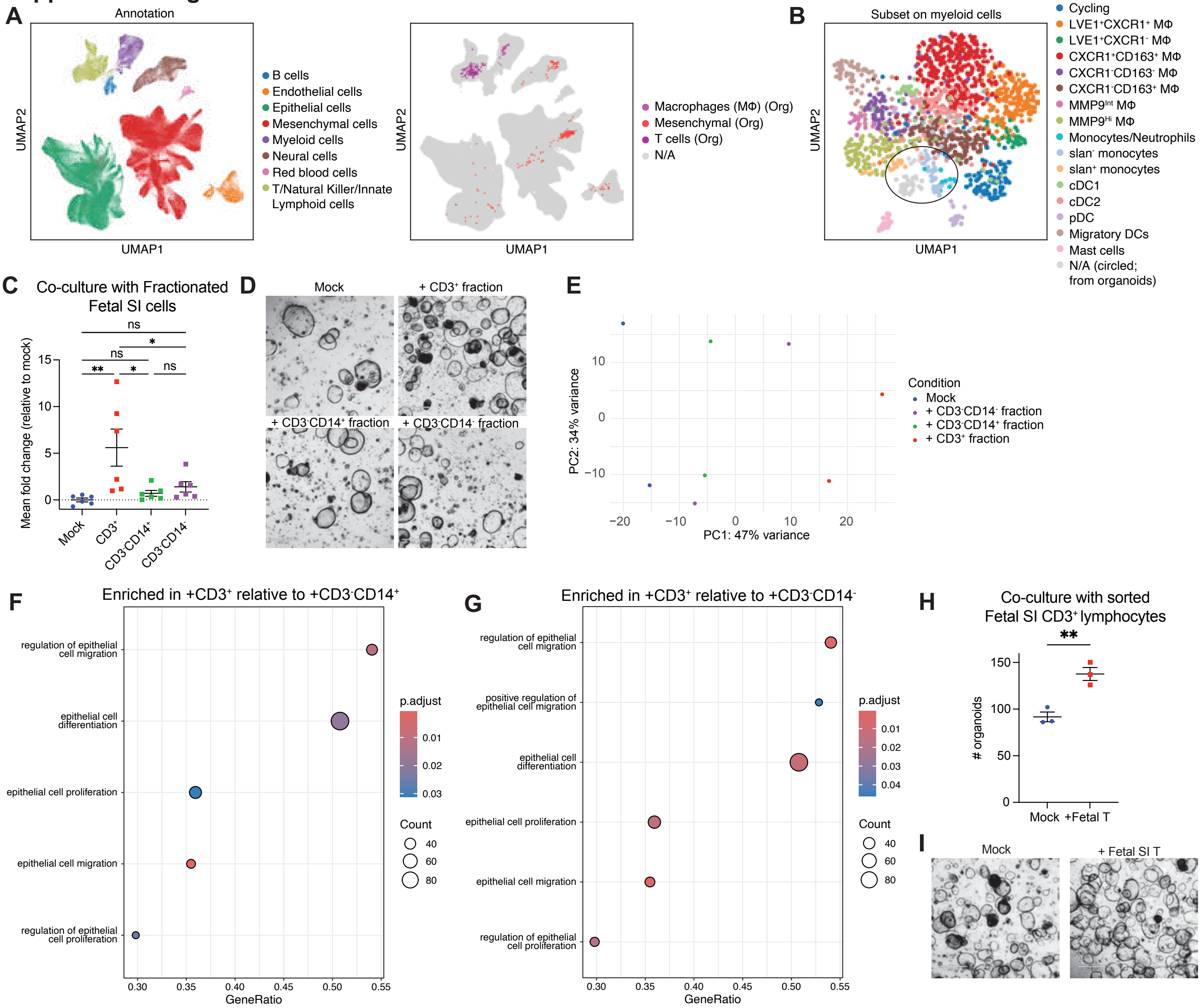

#### Supplemental Figure 5.

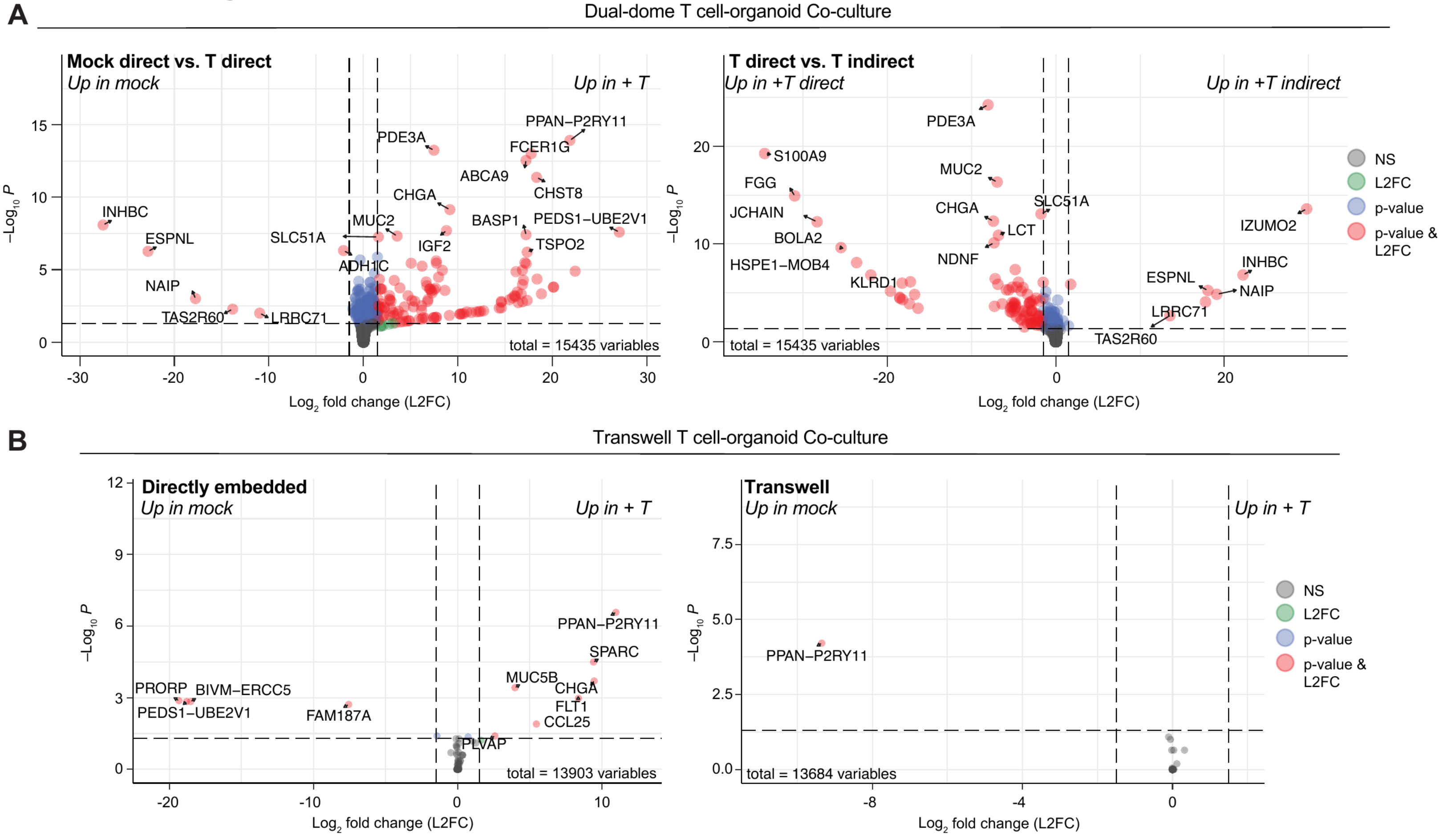

### Supplemental Figure 6.

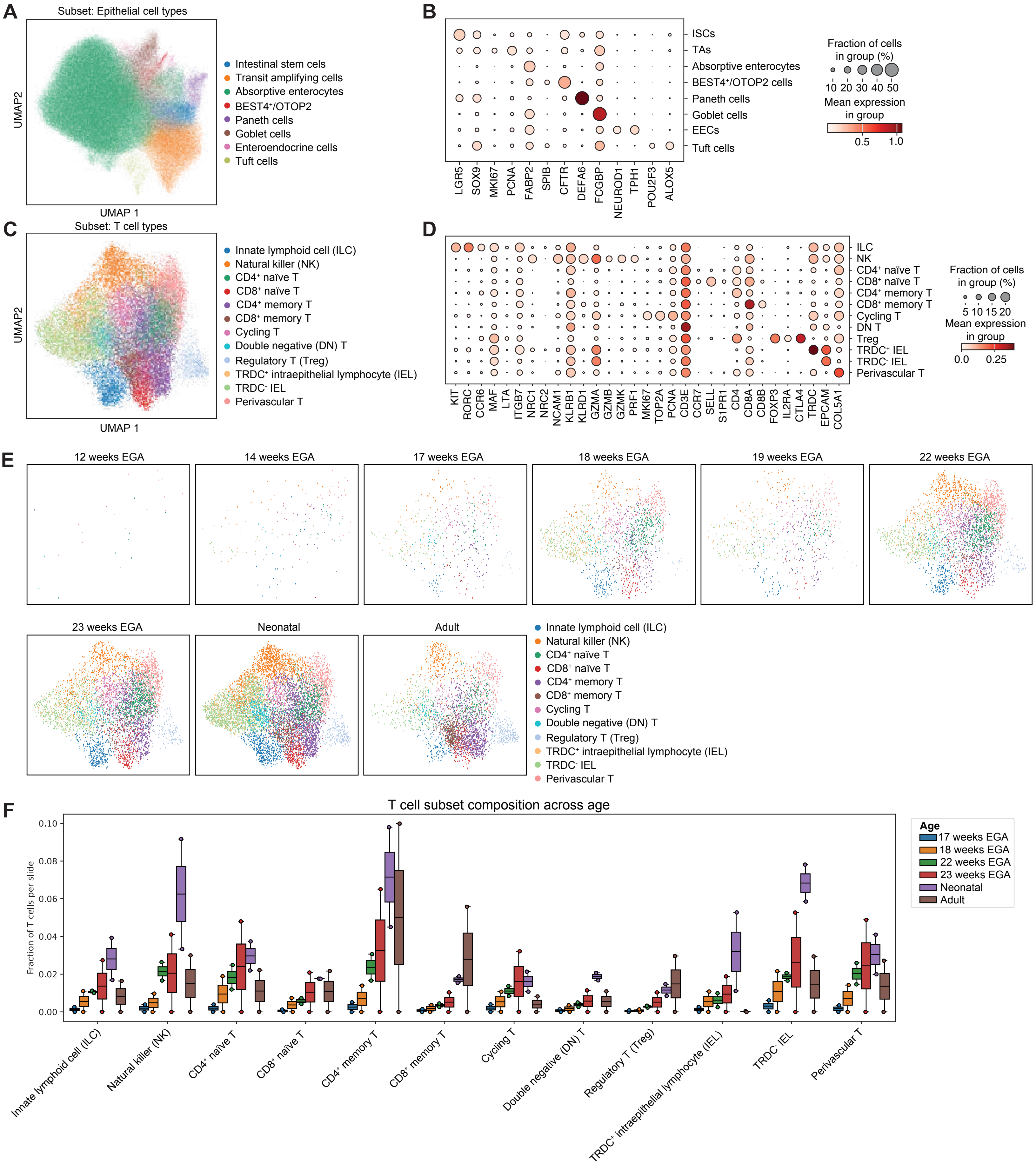

### Supplemental Figure 7.

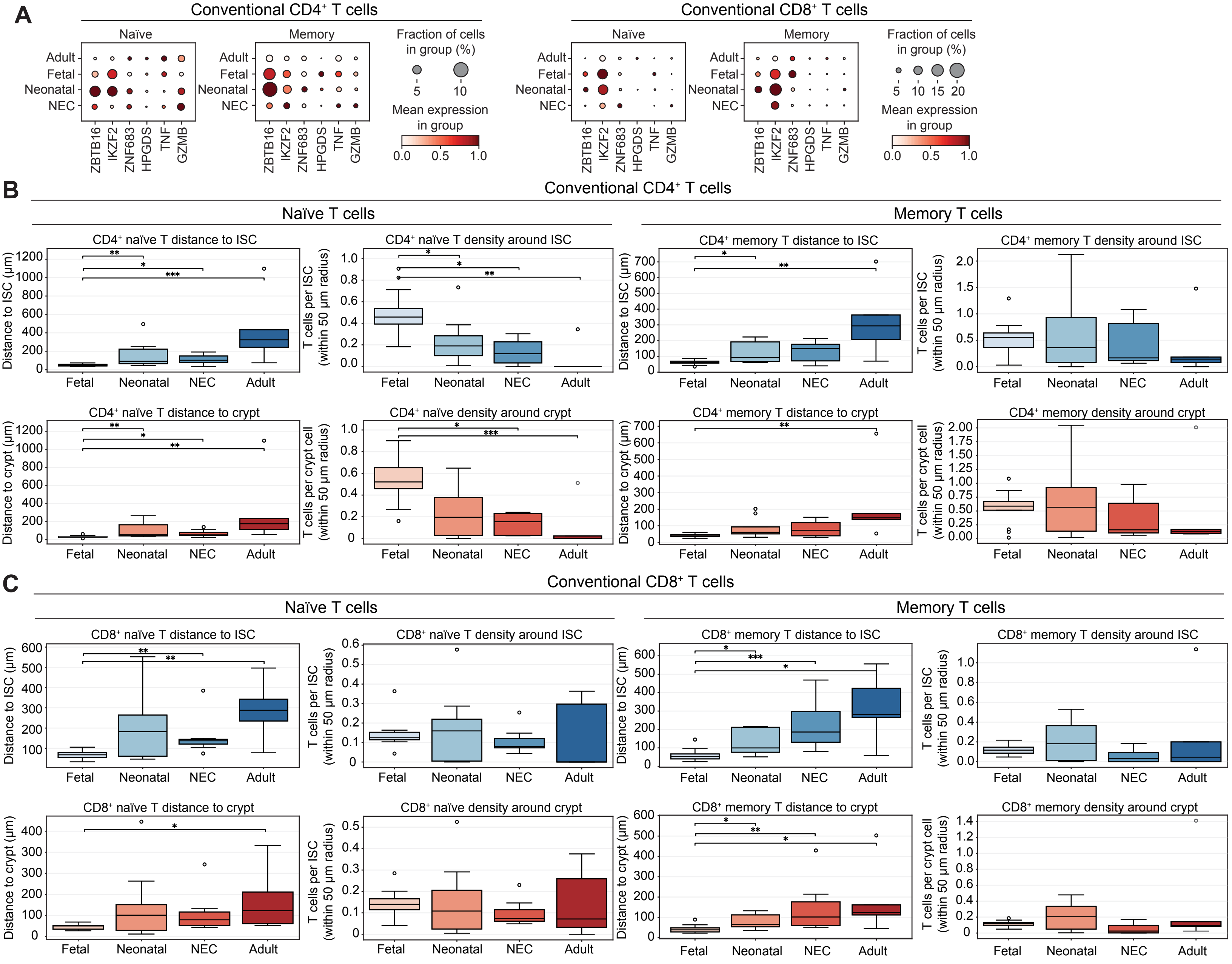
